# Programmable self-assembling system to model intracellular protein aggregation and decode neurodegenerative diseases

**DOI:** 10.1101/2025.03.06.641009

**Authors:** Yujie Fan, Jonathan D. Hoang, Chelsie Steele, Yochana Benchetrit, Sayan Dutta, Gerard M. Coughlin, Nathan Appling, Xiaozhe Ding, Changfan Lin, Zhe Qu, Elisha D. Mackey, Viviana Gradinaru

## Abstract

Pathological protein aggregation is a conserved feature of neurodegenerative diseases. However, slow progression, aggregates heterogeneity and selective vulnerability make it challenging to model diseases and dissect mechanisms. Here, we present a genetically-encoded, modular platform of self-assembling protein that enables inducible, tunable and cell-type-specific formation of intracellular protein aggregates. We engineered self-assembling variants of α-synuclein (SAS), Tau, and TDP-43, recapitulating hallmarks of Parkinson’s disease, Alzheimer’s disease, and ALS, respectively. SAS formed Lewy body-like inclusions, nucleated endogenous α-synuclein, triggered neuroinflammation and neurite degeneration; Self-assembling-Tau induced tangles formation, extracellular Aβ and severe neurodegeneration; Self-assembling-TDP caused TDP-43 mislocalization and cytoplasmic inclusions with mild degeneration. Notably, the system revealed disease-specific aggregate-organelle interactions. *In vivo*, systemic or dopamine neuron-targeted delivery of SAS induced progressive motor deficits, dopaminergic neuron loss, and microglial activation. Our self-assembling system robustly mirrors histological, transcriptional, and behavioral features of human disease, providing a powerful platform to dissect mechanisms and accelerate therapeutic development.

**Graphical abstract:** 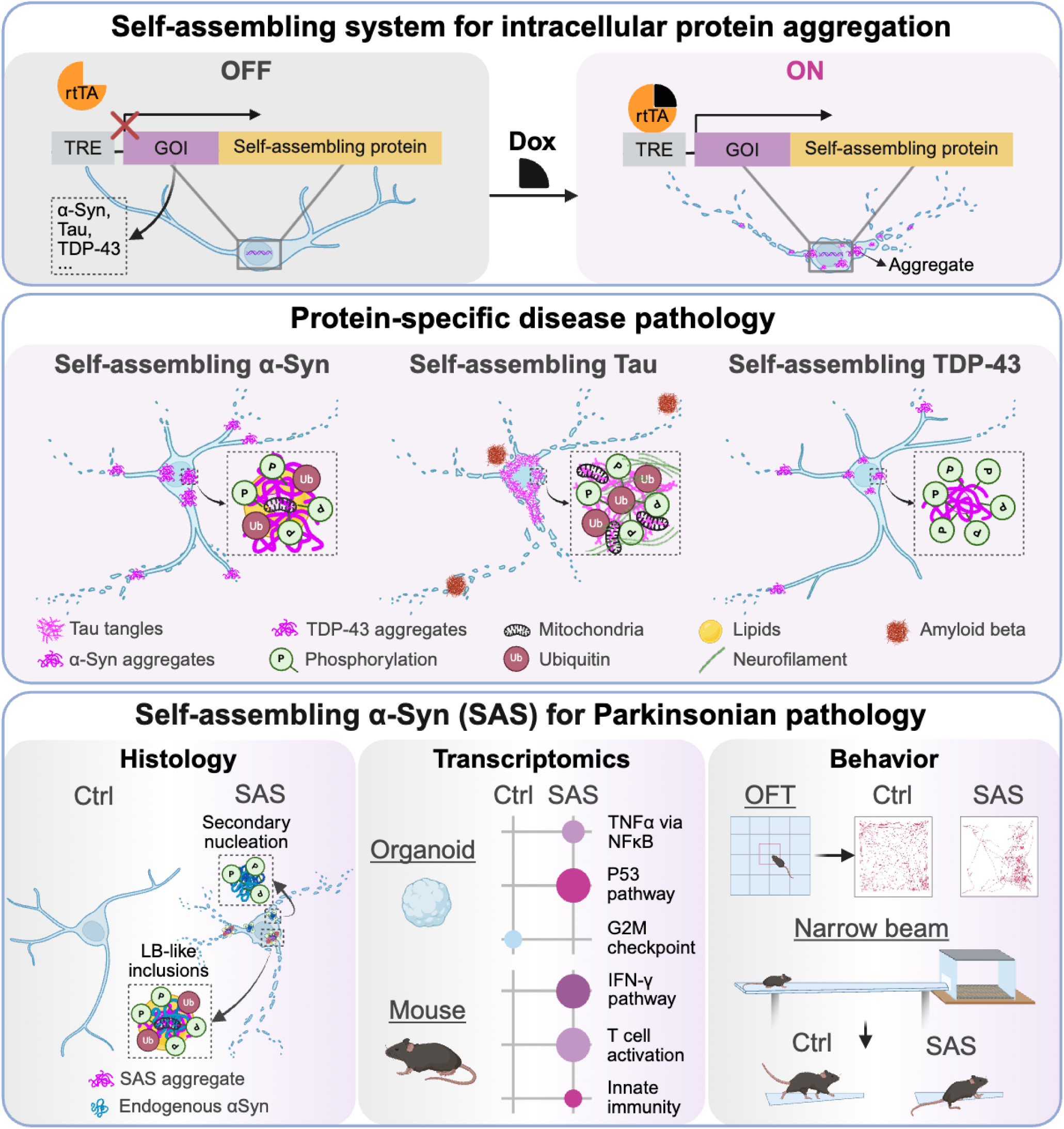

## Introduction

Protein aggregation is a pathological hallmark of numerous neurodegenerative disorders, including Parkinson’s disease (PD), Alzheimer’s disease (AD), and amyotrophic lateral sclerosis (ALS)^1^. In these conditions, misfolded proteins form intracellular or extracellular inclusions that disrupt cellular homeostasis, impair organelle function, propagate through neural circuits and eventually lead to neuronal loss and behavioral changes^2,3^. Despite their shared etiology of protein aggregation, different neurodegenerative diseases involve distinct aggregating proteins, affect different brain regions, exhibit unique patterns of cell-type vulnerability, and lead to divergent clinical outcomes^4^. It remains poorly understood which forms of aggregates are toxic, how aggregates spread between cells and drive neurotoxicity, and why certain neurons are more vulnerable to specific protein aggregates than others.

Protein misfolding and aggregation may begin decades before onset of clinical symptoms and the aggregates display substantial heterogeneity across patients^5^. Capturing the slow progression, heterogeneity and cell-type-specific vulnerability characteristic of human proteinopathies in experimental models remains a major challenge. Most existing models rely on toxicants, overexpression of disease-related proteins or exogenous seeding with fibrils. These approaches often lack temporal and cell-type control and fail to recapitulate key aspects of disease pathology. A notable example is PD, which is characterized by the progressive loss of dopaminergic (DA) neurons and the formation of intracellular inclusions known as Lewy bodies (LBs) and Lewy neurites (LNs)^6–8^. The main component of LBs and LNs is misfolded α-synuclein (αSyn). It remains unclear when and where the αSyn aggregation begins, how it spreads between cells, and why the DA neurons are particularly vulnerable to degeneration^9,10^. Numerous *in vitro* and *in vivo* models have been developed to model and study PD^11–13^. Toxicant-based models, such as 6-hydroxydopamine (6-OHDA), induce rapid DA neuron degeneration but typically fail to produce LBs and LNs pathology^14,15^. Genetic models, including overexpression of αSyn, often do not exhibit key pathological hallmarks ^16^. More recently, the αSyn pre-formed fibril (PFF) model has gained prominence, as intracerebral injection of αSyn PFFs induces neurodegeneration and behavioral deficits in rodents^17–19^. However, this model requires invasive surgical procedures and lacks cell-type specificity. A system that allows precise and programmable modeling of protein aggregation in defined cell types could transform our ability to study disease mechanisms across diverse neurodegenerative conditions.

Here, we developed a genetically-encoded, modular platform that enables inducible, tunable, and cell-type-specific formation of intracellular protein aggregates across diverse *in vitro* and *in vivo* experimental models. Using this platform, we engineered self-assembling variants of αSyn, Tau, and TDP-43 to model hallmark features of PD, AD, and ALS, respectively. In HEK cells, primary neurons and human brain organoids, self-assembling αSyn (SAS), Tau (SA-Tau) and TDP-43 (SA-TDP) induced histopathology that closely mirrored that in the human diseases, establishing a new paradigm in disease modeling. SAS aggregates formed LB-like inclusions that interacted with mitochondria, lipids, lysosomes, and ubiquitin, features observed in PD patients but not fully captured by prior experimental models. Applying the same approach to Tau produced hyperphosphorylated tangles that were highly ubiquitinated, elevated neurofilament levels, and promoted extracellular amyloid beta production, resulting in severe neurodegeneration. In contrast, self-assembling TDP-43 formed phosphorylated cytoplasmic inclusions that exhibited minimal interaction with mitochondria and ubiquitin, leading to milder neurodegeneration at comparable expression levels and time. Furthermore, non-invasive, cell-type-specific delivery of SAS to DA neurons in the mouse brain led to motor deficits, DA neuron loss, reduced striatal projections, and propagation of αSyn aggregates from the substantia nigra pars compacta (SNpc) to other brain regions. This system not only recapitulates key histological and transcriptional features of human disease but also enables programmable modeling of aggregate biology across diverse contexts. Our platform provides a powerful, extensible tool to interrogate mechanisms of neurodegeneration and to accelerate therapeutic development.

## Results

### Self-assembling system induces intracellular protein aggregates and enables tunable proteinopathy

Abnormal protein aggregates in neurodegenerative diseases are composed of not only misfolded protein, but also diverse cellular components^20,21^. Rather than synthesizing pure protein fibrils in solution and then introducing them into cells, we hypothesized that inducing protein aggregation intracellularly would enable the aggregates to incorporate endogenous cellular components, therefore more faithfully recapitulating disease pathology. Previous studies have shown that de novo-designed helical filaments^22,23^ and naturally occurring self-assembling proteins^24,25^ can form ordered aggregates spontaneously in solution or in yeast. Such self-assembling proteins have been applied to record protein activity *in vivo*^26,27^. Inspired by these studies, we developed a genetically encoded platform for programmable aggregate formation, Focusing initially on αSyn, we designed a library of self-assembling synuclein (SAS) constructs, each consisting of human *SNCA*, the gene encoding αSyn, fused with a different self-assembling protein (SAP) at a position predicted to remain accessible on the exterior surface of the resulting filament (**Table S1**). To enable temporal control, we integrated this design into a doxycycline-regulated system. Cell-type specificity was achieved by utilizing custom promoters to drive expression of the reverse tetracycline-controlled transactivator (rtTA), allowing for targeted induction of αSyn aggregation in desired cell populations (**Fig. 1A**).

**Fig. 1.**
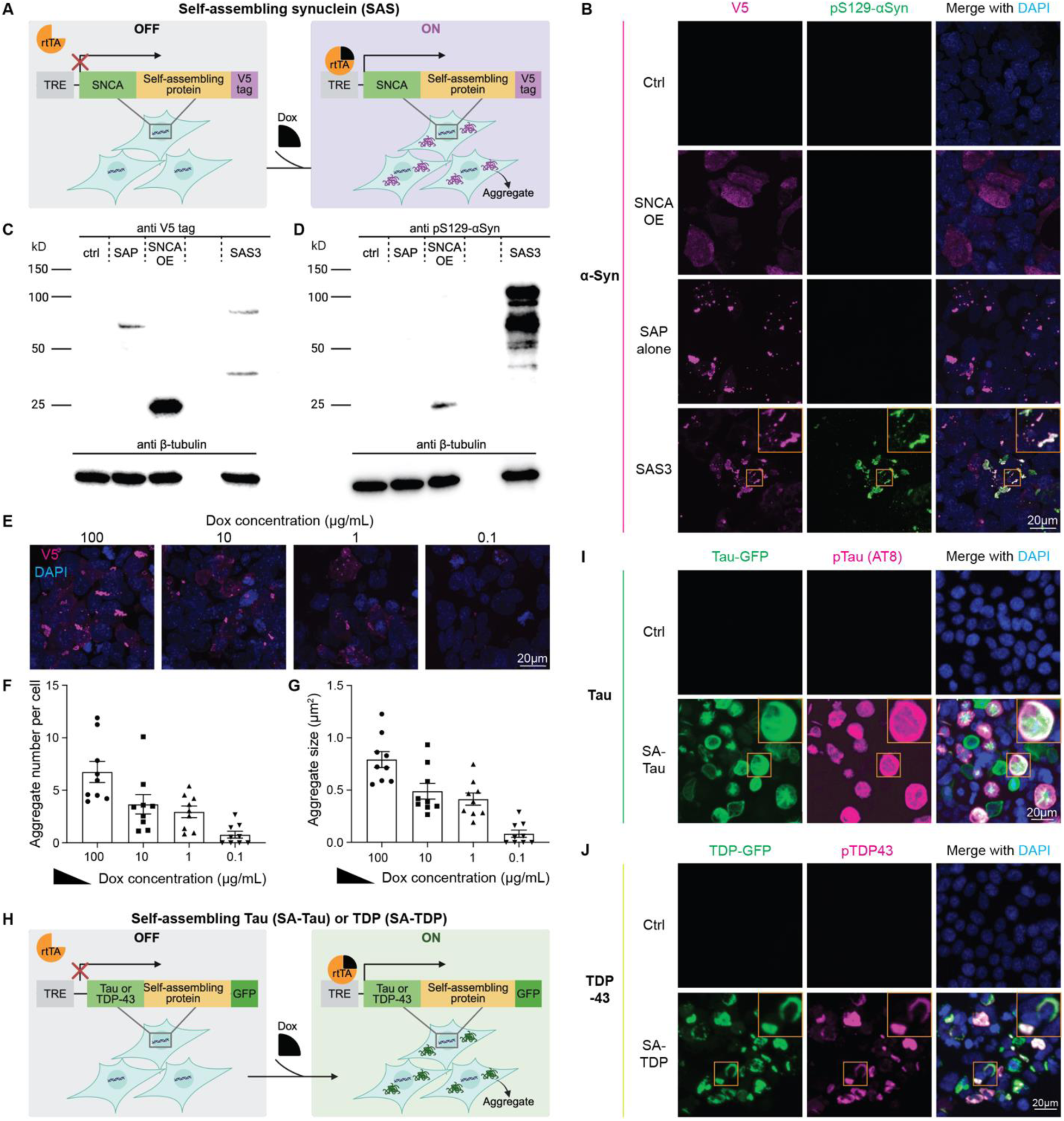
Self-assembling system induces intracellular protein aggregates and enables tunable proteinopathy. **(A)** Design of self-assembling synuclein (SAS) system. SAS consists of αSyn fused with a self-assembling protein. In HEK cells, rtTA expression is driven by the EF1α promoter. Upon doxycycline (dox) induction, SAS forms αSyn aggregates intracellularly. **(B)** Screening of SAS constructs in HEK cells. Compared to ctrl (rtTA-transfected only), *SNCA* overexpression (*SNCA* OE) or self-assembling protein (SAP) alone, SAS3 induces αSyn aggregation and strong αSyn S129 phosphorylation after 3 days of dox treatment in transfected HEK cells. **(C-D)** Western blot analysis of total expression (V5 tag, **C**) and pS129-αSyn (**D**), reveals SAS3-induced aggregates trigger markedly stronger αSyn phosphorylation than *SNCA* OE, suggesting enhanced pathogenic potential. n=3 biological replicates. **e,** Representative images of SAS-induced aggregates at varing dox concentration. **(F-G)** Quantification of aggregate number per cell (**F**) and average aggregate size (**G**) for different dox concentrations, demonstrating tunability of the system. n=9 biological replicates. **(H)** Schematic of modular design for self-assembling Tau (SA-Tau) and self-assembling TDP-43 (SA-TDP), in which human Tau (isoform 2N4R) or TDP-43 is fused with the same self-assembling protein used in SAS3. A GFP tag is used to detect aggregates. **(I)** Immunostaining of HEK cells transduced with SA-Tau-GFP and stained for pTau (AT8) after 3 days of dox induction shows robust Tau aggregation and hyper phosphorylation. **(J)** Immunostaining of SA-TDP-GFP-expressing HEK cells after 3 days of dox induction reveals cytoplasmic TDP-43 inclusions with strong pTDP-43 signal. Scale bars equal 20 μm. Data are presented as mean ± SEM.

*In vitro* screening in HEK cells identified several constructs that not only formed αSyn aggregates but also induced substantial Ser129 phosphorylation of αSyn (pS129-αSyn), a hallmark of PD pathology (**Fig. 1B and S1A**). Different SASs resulted in different types of aggregates with varying ability to induce pS129-αSyn. Among these, SAS3 formed condensed punctate structures and robustly induced pS129-αSyn. To confirm that the pS129-αSyn induction was specific to the αSyn aggregates, we compared SAS3 with rtTA-transfected control (Ctrl), *SNCA* overexpression (*SNCA* OE), and SAP alone. Only SAS3 led to strong pS129 induction; Neither αSyn OE nor SAP alone triggered similar pathology (**Fig. 1B**). Western blot analysis revealed that SAS3 induced 1000-fold higher pS129-αSyn compared to *SNCA* OE (**Fig. 1C&D and S1B&C**), underscoring the pathological potency of the construct. To enable real-time visualization of aggregate dynamics, we generated a fluorescent version of SAS3 by replacing the V5 epitope tag with a GFP reporter (SAS-GFP) (**Fig. S1D&E**). Live imaging demonstrated gradual appearance and growth of intracellular αSyn aggregates (**Movie S1**). The SAS system was designed to allow precise control of αSyn aggregation and pathology through manipulation of doxycycline (dox) concentration and treatment duration. Indeed, titration of dox concentration decreased the average number and size of aggregates (**Fig. 1E, F, and G**). Prolonged dox treatment led to a corresponding increase in aggregation and αSyn phosphorylation (**Fig. S1F**).

To assess the generalizability of the self-assembling protein platform beyond synuclein, we next developed analogous constructs targeting two other major neurodegenerative disease proteins: Tau and TDP-43, which are central to AD and ALS, respectively. Using the same modular design as SAS, we fused Tau (2N4R isoform) or TDP-43 to the self-assembling domains used in SAS3 (**Fig. 1H; Table S1**). In HEK cells, self-assembling Tau (SA-Tau) formed fibrillar cytoplasmic tangles that were positive for phospho-Tau (AT8), a hallmark of tauopathy and AD (**Fig. 1I**). Self-assembling TDP-43 (SA-TDP) induced cytoplasmic inclusions and mislocalization of TDP-43 with strong phospho-TDP-43 signal, mirroring ALS pathology (**Fig. 1J**).

Together, these results demonstrate that our self-assembling system enables intracellular formation of protein aggregates, relevant to multiple neurodegenerative diseases. Its compatibility with genetic control, tunability, and modularity provide robust, scalable, and refined preclinical models for studying disease mechanisms and developing therapeutics.

### Self-assembling αSyn, Tau, and TDP-43 aggregates recapitulate distinct, protein-specific histopathology in neurons

Another defining feature of neurodegenerative diseases is neuronal degeneration. Having shown that our self-assembling system drives robust, phosphorylated protein aggregation in HEK cells, we next investigated whether these aggregates could trigger neuronal degeneration. We delivered the self-assembling system to mouse primary neurons using adeno-associated virus (AAV) AAV9-X1.1^28^, an engineered AAV that efficiently transduces primary murine and human neurons in culture^29^ (**Fig. 2A)**. As observed in HEK cells, SAS3 induced prominent αSyn aggregation and pS129-αSyn in both the soma and neurites of primary neurons (**Fig. 2B** and quantified in **Fig. 2C&D**). Notably, SAS3-transduced neurons also exhibited pronounced neurite retraction after 3 days of dox treatment (**Fig. 2B&E**), mimicking the neurodegeneration observed in PD. Consistent with these findings, SAS3 also induced αSyn aggregation and neurite retraction in human embryonic stem cell (hESC)-derived neurons (**Fig. S2A**). Similarly, SA-Tau formed extensive tau tangles positive for phospho-Tau (AT8), accompanied by severe neuronal loss and neurite degeneration as marked by MAP2 (**Fig. 2F**). Intriguingly, SA-Tau expression also led to elevated levels of extracellular amyloid beta (Aβ) (**Fig. 2G**), suggesting a potential link between intracellular Tau aggregation and Aβ dysregulation. Neurons expressing SA-TDP exhibited cytoplasmic TDP-43 inclusions with strong pTDP-43. However, compared with equivalent expression of SAS and SA-Tau, SA-TDP induced milder neurodegeneration (**Fig. 2H**). Total TDP-43 staining revealed translocation of endogenous TDP-43 from the nucleus to cytoplasm and aggregation (**Fig. 2I**). Collectively, these findings establish that the self-assembling protein aggregates not only induce neurodegeneration, but also recapitulate distinct, protein-specific pathological signatures.

**Fig. 2.**
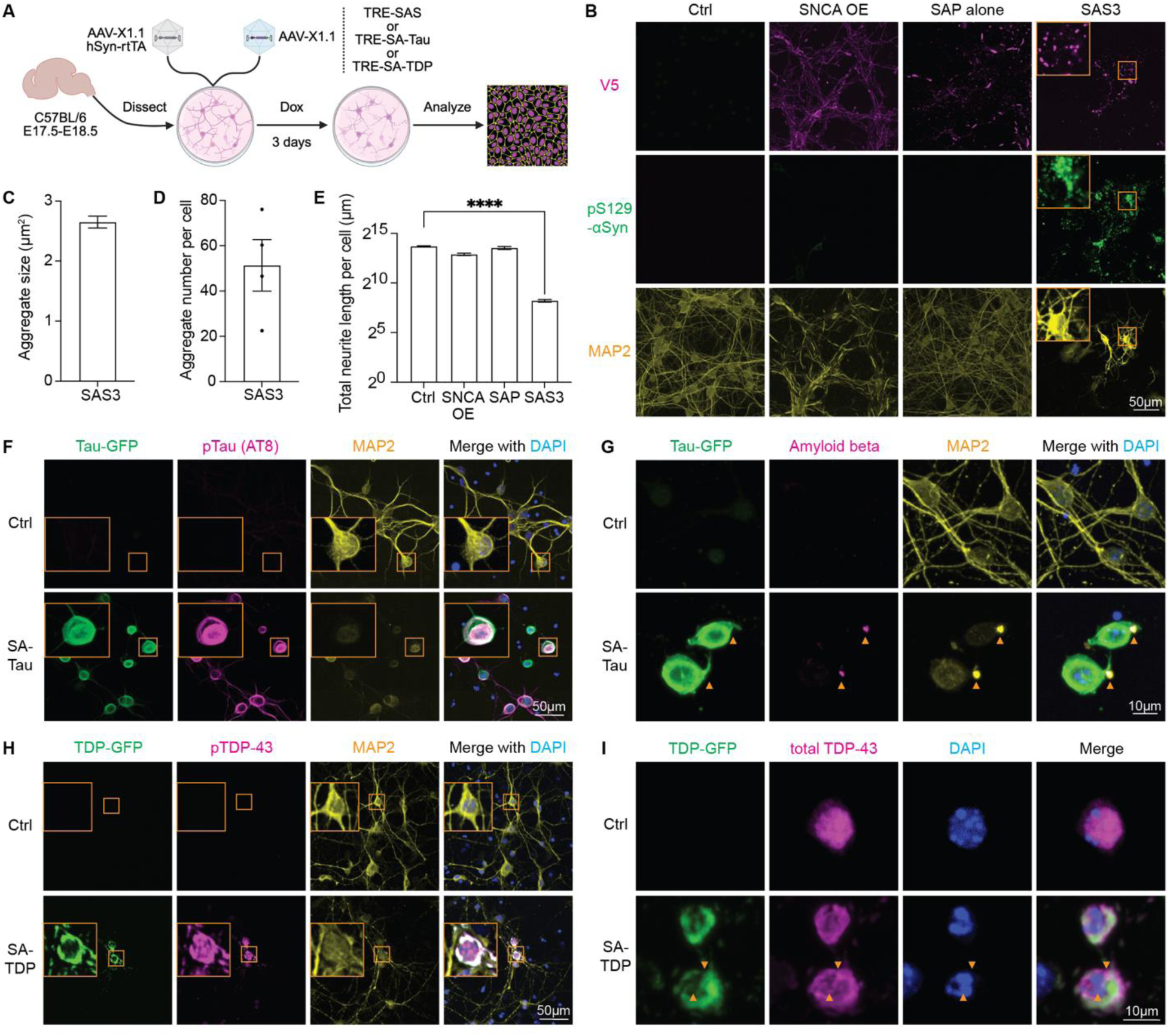
Self-assembling αSyn, Tau, and TDP-43 aggregates recapitulate distinct, protein-specific histopathology in neurons. **(A)** Experimental design for testing the self-assembling system in mouse primary neurons, which is delivered by AAV9-X1.1. In primary neurons, rtTA expression is driven by the hSyn promoter. **(B)** Immunostaining of V5 tag, pS129-αSyn, and MAP2 in transduced primary neurons after 3 days of dox treatment, showing robust αSyn aggregation and pS129-αSyn in both the soma and neurites of neurons, as well as neurite retraction. **(C)** Quantification of average aggregate size induced by SAS3 in mouse primary neurons. **(D)** Average number of aggregates per cell. **(E)** Quantification of total neurite length per cell. n=4 biological replicates. ****p<0.0001. **(F)** Immunostaining of GFP, pTau (AT8 antibody), and MAP2 in SA-Tau-transduced neurons after 3 days of dox treatment, reveals robust Tau aggregation, hyper pTau, and pronounced neurite degeneration. Enlarged images demonstrate that SA-Tau and SASs form distinct aggregate structures, despite the use of the same self-assembling protein. **(G)** Immunostaining for amyloid beta in primary neurons expressing SA-Tau indicates Tau tangles promote the production of extracellular amyloid beta (arrowhead). **(H)** Representative images of neurons expressing SA-TDP after 3 days of dox treatment show cytoplasmic TDP-43 inclusions with strong pTDP-43. Compared with equivalent expression of SAS and SA-Tau, SA-TDP induces milder neurodegeneration. **(I)** Immunostaining for total TDP-43 reveals translocation of endogenous TDP-43 from nucleus to cytoplasm and aggregation (arrowhead). Scale bar equals 50 μm in **B**, **F, H,** and 10 μm in **G, I**. Data are presented as mean ± SEM; statistical analysis was performed using ANOVA with Dunnett’s test.

### Self-assembling system reveals distinct organelle interactions and pathological signatures of αSyn, Tau, and TDP-43 aggregates in human brain organoids

How different protein aggregates drive distinct neurotoxicity remains poorly understood. We hypothesized that differential interactions between protein aggregates and other cellular components may underlie their varying toxic effects, contributing to the pathogenesis of different neurodegenerative diseases^30^. Supporting this idea, postmortem examination of PD patients has shown that aggregates are far from homogeneous lumps of toxic protein^31^. LBs often contain hundreds of different proteins and even entire membranous organelles tangled up in them^32,33^. One considerable advantage of the self-assembling system is its ability to drive intracellular aggregation, potentially enabling incorporation of other cellular components and better recapitulation of the molecular complexity of pathological inclusions.

To test this hypothesis in a physiologically relevant context, we delivered SAS, SA-Tau and SA-TDP43 to hESC-derived brain organoids using AAV-X1.1 (**Fig. 3A**). Organoids produce diverse cell types in a 3D environment that closely mimics the function and structure of *in vivo* counterparts. While organoids have been widely used for modeling neurodevelopment, existing organoid models often fail to display hallmark features of neurodegenerative diseases, such as LB formation and neuronal degeneration in PD^34^. In particular, αSyn PFFs, which have shown promise in animal studies, are inefficient in organoids due to limited uptake and poor tissue penetration (**Fig. 3B**). In contrast, AAV-delivered SAS3 was able to efficiently access the entire organoids. Immunostaining for the V5 tag and pS129-αSyn revealed widespread synucleinopathy, with aggregates present even in deep layers of organoids (**Fig. 3B and S3A&B**). Similarly, SA-Tau and SA-TDP produced hallmark pathologies of AD and ALS in human brain organoids, including phosphorated Tau tangles and TDP-43 aggregates, extracellular Aβ accumulation, and cytoplasmic TDP-43 inclusions (**Fig. S3D-G**). Together, these findings establish the self-assembling system as a robust platform for modeling and investigating authentic inclusion biology in a human-relevant system, filling a major gap in current disease models.

**Fig. 3.**
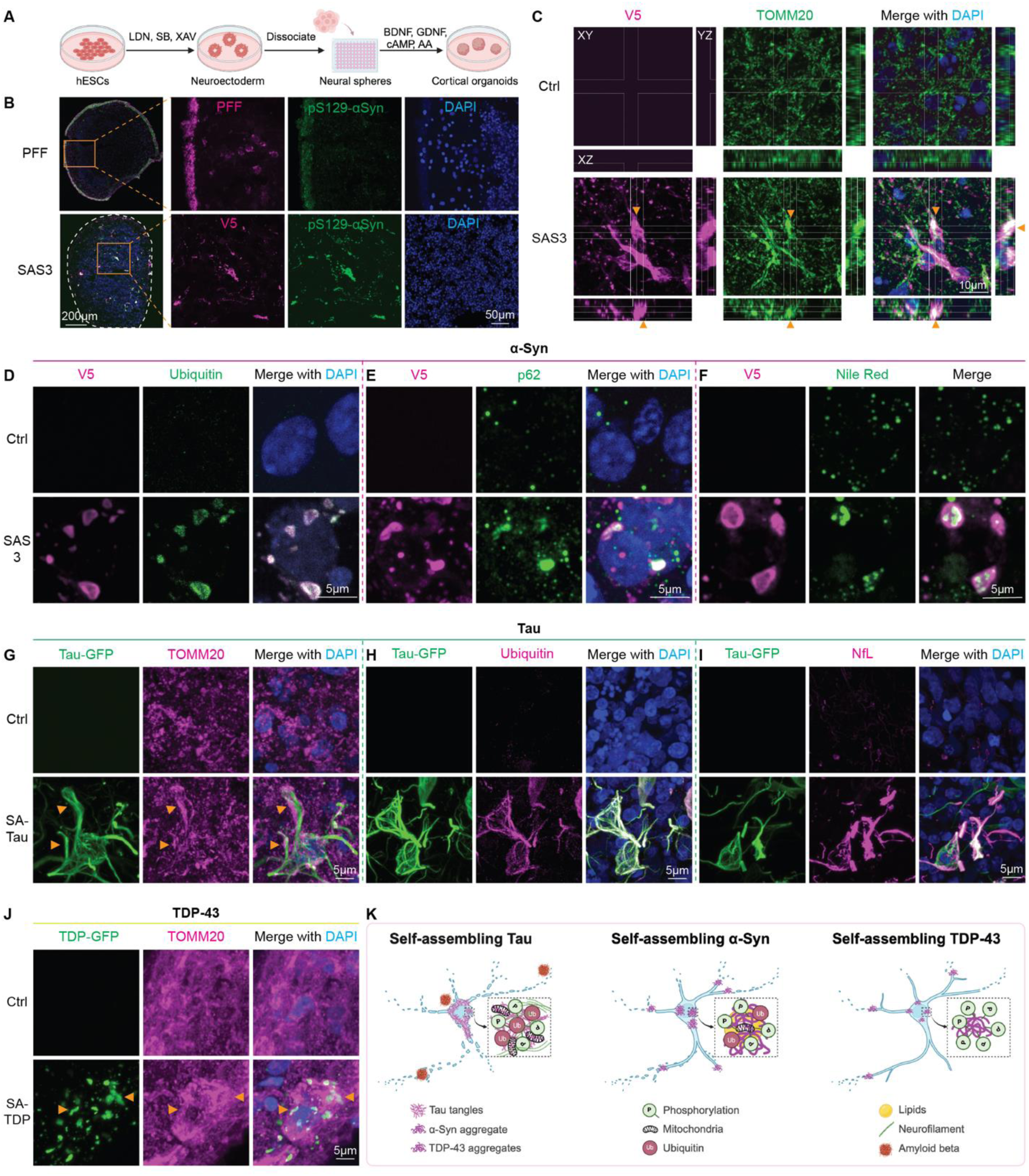
Self-assembling system reveals distinct organelle interactions and pathological signatures of αSyn, Tau, and TDP-43 aggregates in human brain organoids. **(A)** Schematic of generating human cortical organoids from hESCs. **(B)** Immunostaining of fluorescently-labeled PFFs or SAS3 (V5 tag) and pS129-αSyn in hESC-derived cortical organoids. Unlike PFFs, SAS3 can penetrate and induce PD-like pathology through the entire organoid. Scale bar equals 200 μm and 50 μm in enlargement. **(C)** Z-scanning confocal imaging of organoids stained for V5 tag and the mitochondrial marker TOMM20. SAS3-induced aggregates colocalize with mitochondria (arrowhead), shown by XY (main panel), XZ (bottom panel) and YZ (right panel) projections. Scale bar equals 10 μm. **(D-F)** Organoids stained for ubiquitin and p62/SQSTM1 or with Nile red reveal that SAS3-induced aggregates are complex inclusions containing αSyn and other proteins, similar to LBs found in human PD patients. **(D)** Ubiquitin staining confirms that proteins in SAS3-induced aggregates are highly ubiquitinated. **(E)** Accumulation of p62 in and around the aggregates indicates elevated autophagy. **(F)** Nile red staining demonstrates enrichment of lipids in SAS-induced aggregates. **(G)** Mitochondrial marker TOMM20 accumulates surrounding SA-Tau aggregates (arrowhead). **(H-I)** SA-Tau-induced aggregates are highly ubiquitinated (**H**) and co-localize with neurofilament light chain (NfL) (**I**). **(J)** In contrast, SA-TDP-induced aggregates show little to no co-localization with mitochondria (arrowhead), indicating a distinct aggregate composition compared to SAS and SA-Tau. **(K)** Schematic comparison of SAS, SA-Tau and SA-TDP aggregates, highlighting differences in aggregate morphology, interaction patterns with organelles, and associated neurotoxicity. Scale bar equals 5 μm in **D-J**.

To investigate the molecular composition and cellular impact of aggregates, we performed confocal Z-stack imaging of organoids infected with self-assembling constructs and stained for cellular components. In SAS-expressing organoids, co-staining for the V5 tag to trace aggregates and the mitochondrial membrane marker TOMM20 revealed clear TOMM20 signal within the αSyn aggregates, indicating that mitochondria were sequestered in the inclusions (**Fig. 3C**), a feature not observed in control groups (**Fig. S3C**). SAS-induced inclusions were highly ubiquitinated (**Fig. 3D**) and showed prominent accumulation of autophagy adaptor p62/SQSTM1 in and around the aggregates (**Fig. 3E**), suggesting these inclusions are recognized and targeted by protein clearance pathways. Nile red staining revealed that aggregates were enriched in lipids (**Fig. 3F**), a common feature of LBs from PD patient brains. Tau and TDP-43 aggregates displayed distinct aggregate-organelle interaction profiles. In SA-Tau-expressing organoids, mitochondria accumulated around the aggregates rather than being trapped inside (**Fig. 3G**). Tau tangles were also highly ubiquitinated (**Fig. 3H**), p62-positive (**Fig. S3F**) and colocalized with neurofilament light chain (NfL) **(Fig. 3I)**. In contrast, SA-TDP-induced aggregates showed minimal co-localization with mitochondria, ubiquitin, or NfL, but still induced p62 accumulation (**Fig. 3J and S3H-J**). This divergence in aggregate-organelle interactions (**Fig. 3K**) may explain the milder neurodegeneration observed in SA-TDP-expressing organoids and primary neurons compared to SAS and SA-Tau.

Self-assembling protein system recreated protein-specific aggregation and disease-relevant pathology in human brain organoids and uncovered distinct aggregate-organelle interaction profiles that might underlie differential aggregates toxicity. The ability to resolve these aggregate-specific signatures provides a unique opportunity to dissect the molecular basis of selective vulnerability in PD, AD, ALS, and other related disorders.

### SAS forms LB-like inclusions and nucleate aggregation of endogenous αSyn, triggering neuronal inflammation and death

Having established our self-assembling system faithfully recapitulates protein-specific pathology across neurodegenerative diseases, we next focused on PD to further explore the full potential of SAS in elucidating mechanisms of αSyn-mediated neurodegeneration. Our findings demonstrated that SAS induced complex, multi-component inclusions that closely resemble human LBs in terms of molecular composition^38^ (**Fig. 3C-F**). Here, we further showed the inclusion also recapitulate key morphological features of LBs. Thioflavin S staining confirmed the presence of β-sheet–rich fibrillar content in SAS-expressing neurons (**Fig. S4A**). Notably, the Thioflavin S signal often extended beyond the V5-positive core, suggesting the existence of secondary aggregated species. Co-staining of the V5 tag and pS129-αSyn revealed numerous V5-negative and pS129-αSyn-positive aggregates, pointing to the formation of endogenous αSyn inclusions (**Fig. S4B**). PD is widely considered a prion-like disease, in which abnormal αSyn aggregates can induce secondary nucleation of normal αSyn^35–37^, promoting disease spreading. To determine whether SAS-induced aggregates possess this capability, we used primary neurons carrying an SNCA-GFP reporter, enabling tracking of the endogenous αSyn (**Fig. 4A**). Following 36 hours of dox treatment, neurons expressing SAS3 exhibited degeneration phenotypes compared with control groups (**Fig. 4B**), with further worsening by 72 hours (**Fig. S4C**). Close examination of SAS3-transduced neurons revealed endogenous αSyn aggregation in both soma and neurites (**Fig. 4C**), confirming a prion-like seeding effect.

**Fig. 4.**
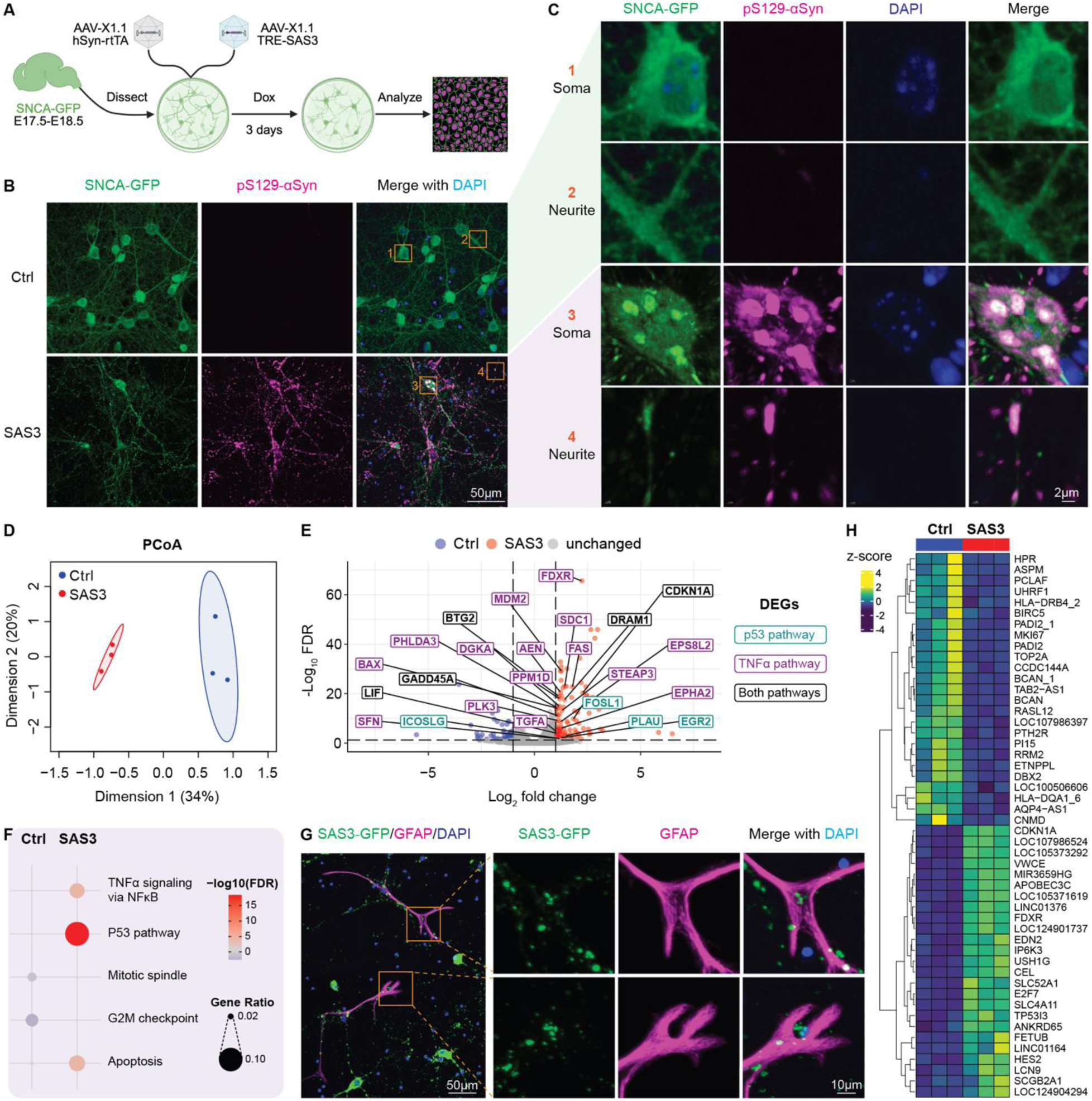
SAS forms LB-like inclusions and nucleate aggregation of endogenous αSyn, triggering neuronal inflammation and death. **(A)** Experimental design for testing whether SAS3 can nucleate secondary aggregation of endogenous αSyn in primary neurons carrying the SNCA-GFP reporter. **(B-C)** Immunostaining after 36 hours of dox treatment shows SAS3-induced aggregates contain endogenous αSyn, as indicated by GFP signal. Scale bar equals 50 μm in (**B**) and 2 μm in (**C**). **(D)** Principal coordinate analysis (PCoA) shows a clear separation of transcriptomic profiles of SAS3 organoids transduced with hSyn-rtTA and TRE-SAS3 using AAV9-X1.1 and control organoids transduced with only hSyn-rtTA. n= 3 biological replicates. **(E)** Volcano plot of DEGs with FDR < 0.05 and |log_2_ fold change (FC)| > 1. Genes belonging to the TNFα pathway are highlighted in magenta, p53 pathway in cyan, and both in black. **(F)** Pathway analysis reveals that SAS3-treated organoids contain significantly upregulated Gene Ontology (GO) pathways associated with apoptosis and inflammation. **(G)** Immunostaining for the astrocyte marker GFAP reveals astrocyte activation surrounding SAS3-expressing neurons and active engagement with protein inclusions. Scale bar equals 50 µm in overview and 10 µm in enlargement. **(H)** Top 25 up- and downregulated genes.

To better understand the mechanisms of SAS-induced neurodegeneration, we conducted bulk RNAseq of SAS-transduced and control organoids. SAS-expressing organoids displayed a distinct transcriptomic profile (**Fig. 4D**), with 178 differentially expressed genes (DEGs) with adjusted p-value < 0.05 and |log_2_ fold change (FC)| >1 (**Fig. S4D** and **Table S2**). Pathway analysis revealed significant upregulation of p53 and TNFα signaling, key regulators of apoptosis and inflammation (**Fig. 4E**). Notably, the TNF-NF-κB-p53 axis has been demonstrated to mediate dopamine neuron survival in neuron transplant experiments^39^. Targeting this pathway might enhance neuron survival in PD. Consistent with human transcriptomic data^40^, SAS-treated organoids showed activation of apoptosis, inflammation, and mitochondrial dysfunction pathways (**Fig. 4F and S4E**). In line with transcriptomic findings, immunostaining for the astrocyte marker GFAP revealed astrocyte activation surrounding SAS-expression neurons and active engagement with protein inclusions (**Fig. 4G**). Among the top 50 most dysregulated genes, many were previously implicated in PD (**Fig. 4H**). For example, *FDXR*, which encodes the mitochondrial ferredoxin reductase that helps produce iron-sulfur clusters, has been linked to inflammation-associated neurodegeneration^41^, consistent with our earlier finding that SAS inclusions sequester mitochondria (**Fig. 3C**).

These results illustrated that SAS induced both primary and secondary αSyn aggregation, forming inclusions that resemble human LBs in molecular composition, morphology and pathogenic function. Further, these inclusions triggered TNFα-mediated neuroinflammation, glial activation and ultimately neuronal death.

### Brain-wide delivery of SAS induces PD-relevant histopathology and behavioral deficits in mice

The primary clinical manifestation of PD is movement disorder. To investigate whether SAS can induce behavior abnormalities *in vivo*, we utilized AAV-PHP.eB, an AAV capable of crossing the blood-brain barrier (BBB) after systemic injection, to achieve non-invasive delivery of SAS into the mouse central nervous system (CNS)^42^. To specifically target neurons, we used the human synapsin (hSyn) promoter to drive the expression of rtTA. The hSyn-rtTA and TRE-SAS3 constructs were packaged into AAV-PHP.eB and co-delivered into wild-type (WT) C57BL/6J mice via retro-orbital (RO) injection. Dox treatment started 2 weeks after virus injection and open field test (OFT) assay were performed every other week (**Fig. 5A**). After 4 weeks of dox treatment, SAS3-treated mice exhibited a progressive decline in movement in the OFT compared to control groups (**Fig. 5B**). After 8 weeks, mice were sacrificed for histological analysis to assess neuropathological changes associated with SAS3 treatment. We first examined the substantia nigra (SN) and observed aggregates in both SN pars compacta (SNpc) and reticulata (SNr). Strikingly, the SNpc exhibited a clear reduction in TH+ neurons (**Fig. 5C**), and the intensity of DA neuron projections was significantly diminished in the striatum (**Fig. 5D&E**). Next, we examined the olfactory bulb (OB), which PHP.eB strongly transduces. Consistent with our *in vitro* findings, SAS-induced aggregates were associated with robust pS129-αSyn staining, whereas the control (hSyn-rtTA only), *SNCA* OE, and SAP treatments did not induce significant pS129-αSyn signal (**Fig. 5F and S5A**). Aggregates were also found in other brain regions such as cortex and hippocampus (**Fig. S5B** and see **Movie S2** for aggregate distribution in a cleared whole brain). Finally, proteinase K treatment of a SAS3-transduced brain confirmed the resistance of the aggregates to proteolytic degradation (**Fig. 5G**), further supporting their pathological relevance.

**Fig. 5.**
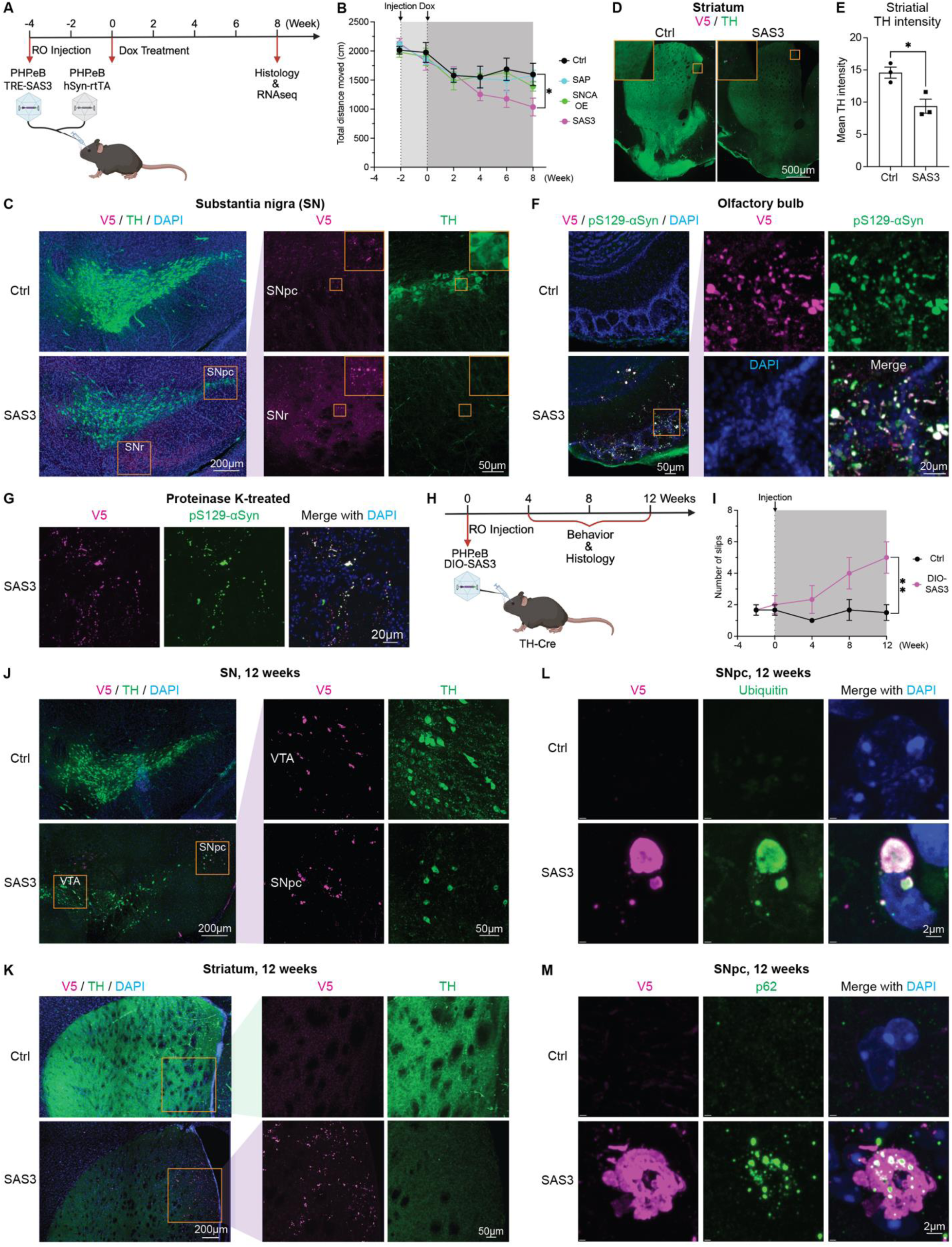
SAS induces PD-relevant histopathology and behavioral deficits in mice. **(A)** Experimental design for brain-wide delivery of SAS3 in WT C57BL/6J mice through retro-orbital (RO) injection of AAV-PHP.eB. Behavioral assays were performed every other week and histology at week 8. **(B)** After 4 weeks of dox treatment, SAS3-treated mice exhibited progressively decreased movement in the open field test compared with other groups. n=5 mice per group. **(C)** Immunostaining of the V5 tag and TH in the substantia nigra, showing aggregates in both the substantia nigra pars compacta (SNpc) and substantia nigra reticulata (SNr), and a decreased number of TH+ neurons in the SNpc. Scale bar equals 200 μm and 50 μm in enlargements. **(D)** Representative images of dopaminergic neuron projections into the striatum. Scale bar equals 500 μm. **(E)** Quantification of striatal TH intensity. n=3 mice. **(F)** Immunostaining of the V5 tag and pS129-αSyn in the olfactory bulb, showing aggregates and strong αSyn phosphorylation. Scale bar equals 50 μm and 20 μm in enlargement. **(G)** Immunostaining for the V5 tag and pS129-αSyn in the olfactory bulb of SAS3-treated mice after proteinase K treatment, showing resistance of aggregates to degradation. Scale bar equals 20 μm. **(H)** Experimental design for selectively delivering SAS3 into dopaminergic neurons. Cre-dependent SAS3 (DIO-SAS3) was RO injected into TH-Cre mice using AAV-PHP.eB. Behavioral assays and histology were performed at 4, 8, and 12 weeks. **(I)** After 4 weeks of dox treatment, SAS3-treated mice exhibited significant motor impairments in the beam crossing assay, displaying more slips than controls. n=5 mice per group. **(J)** Immunostaining of the V5 tag and TH in the substantia nigra (SN) region, with enlarged images of the ventral tegmental area (VTA) and SNpc. **(K)** Immunostaining of the V5 tag and TH in the striatum, suggesting progressive accumulation of aggregates and potential spreading along the nigrostriatal pathway. Scale bar equals 200 μm and 50 μm in enlargements in **J**, **K**. **(L-M)** Immunostaining of ubiquitin **(L)** and p62/SQSTM1 **(M)** confirms that SAS3-induced aggregates are similar to LBs both in morphology and composition. Scale bars equal 2 μm. Data are presented as mean ± SEM; statistical analysis was performed using the ANNOVA with multiple comparison correction post-hoc tests in (**B**) and unpaired t test in (**E&I**); *p<0.05, **p<0.01.

### Dopaminergic neuron-targeted SAS recapitulates progressive motor dysfunction and αSyn aggregate propagation in mice

As a genetically encoded system, SAS provides a well-defined platform to dissect the roles of specific cell types in PD pathology. Degeneration of DA neurons in the SNpc causes motor symptoms in PD^43^. To selectively deliver SAS3 into DA neurons, we injected Cre-dependent SAS3 (DIO-SAS3) into TH-Cre mice using AAV-PHP.eB (**Fig. 5H**). 4 weeks post virus injection, SAS3-treated mice started to exhibit significant motor impairments in the beam-crossing assay, requiring more time to traverse the beam (**Fig. S6A**) and displaying an increased number of slips (**Fig. 5I**) compared to control mice. The motor deficits progressively worsened with prolonged dox treatment, indicating ongoing neurodegeneration. Mice were sacrificed at 4, 8 and 12 weeks for histological analysis to assess the progression of pathology. We first examined DA neurons in the SN (**Fig. 5J and S6B**). At week 4, SAS-induced aggregates were primarily detected in DA neurons within the ventral tegmental area (VTA) and SNpc. By week 8, the number of aggregates had increased, and many TH+ neurons in the SNpc exhibited a clear reduction in TH expression (**Fig. S6B**). At week 12, while most neurons with aggregates in the VTA retained TH expression, a significant proportion of neurons in the SNpc with aggregates exhibited markedly reduced or absent TH expression (**Fig. 5J**). We also detected SAS-induced aggregates in regions that are not typically targeted by TH-Cre, suggesting potential propagation of aggregates beyond DA neurons. A similar trend was observed in the striatum, the primary target of DA neurons from the SNpc. At week 4, no aggregates were detectable in the striatum (**Figure S6C**). However, aggregates emerged by week 8 (**Fig. S6C**) and became more prominent by week 12 (**Fig. 5K**), suggesting progressive aggregate accumulation and potential spreading along the nigrostriatal pathway. The intensity of TH staining in the striatum also decreased with time (**Fig. 5K and S6C**), reflecting a reduction in dopaminergic innervation, consistent with the observed neurodegeneration in the SNpc. Immunostaining for V5 and pS129-αSyn revealed that not only were SAS-induced aggregates strongly phosphorylated (**Fig. S6D, arrows**), but a significant amount of endogenous α-synuclein also aggregated and became phosphorylated (**Fig. S6D, arrowheads**). This again confirmed that SAS3 can induce secondary nucleation of αSyn, contributing to the spread of disease. Ubiquitin and p62/SQSTM1 immunostaining confirmed the molecular complexity of the SAS-induced aggregates and their resemblance to LBs **(Fig. 5L&M)**.

Targeting SAS expression to DA neurons recapitulated multiple hallmarks of Parkinson’s disease, from motor impairment and dopaminergic degeneration to αSyn propagation and LB-like inclusion formation. These results establish SAS as a mechanistically informative platform for probing selective vulnerability and intercellular aggregate dynamics *in vivo*.

### SAS activates microglia and triggers immune response in mice

To dissect transcriptional changes in response to SAS-induced aggregates *in vivo*, we performed bulk RNAseq of the OB, a region strongly transduced by PHP.eB and one of the earliest and consistently affected brain regions in PD. SAS-transduced mice displayed a distinct transcriptome compared with control mice expressing only rtTA (**Fig. 6A**). Differential gene expression analysis identified 139 DEGs with adjusted p-values < 0.05 and |log_2_FC| >1 (**Table S3**), with a strong enrichment of genes promoting inflammation (**Fig. 6B**). Pathway analysis confirmed robust activation of inflammatory and immune responses (**Fig. 6C and S7A**). Among the top 50 most dysregulated genes (**Fig. 6D**), several key inflammatory pathways were highly upregulated, including interferon-gamma (IFN-γ), tumor necrosis factor-alpha (TNF-α), toll-like receptor (TLR), and interleukin (IL) signaling (**Fig. 6E**).

**Fig 6:**
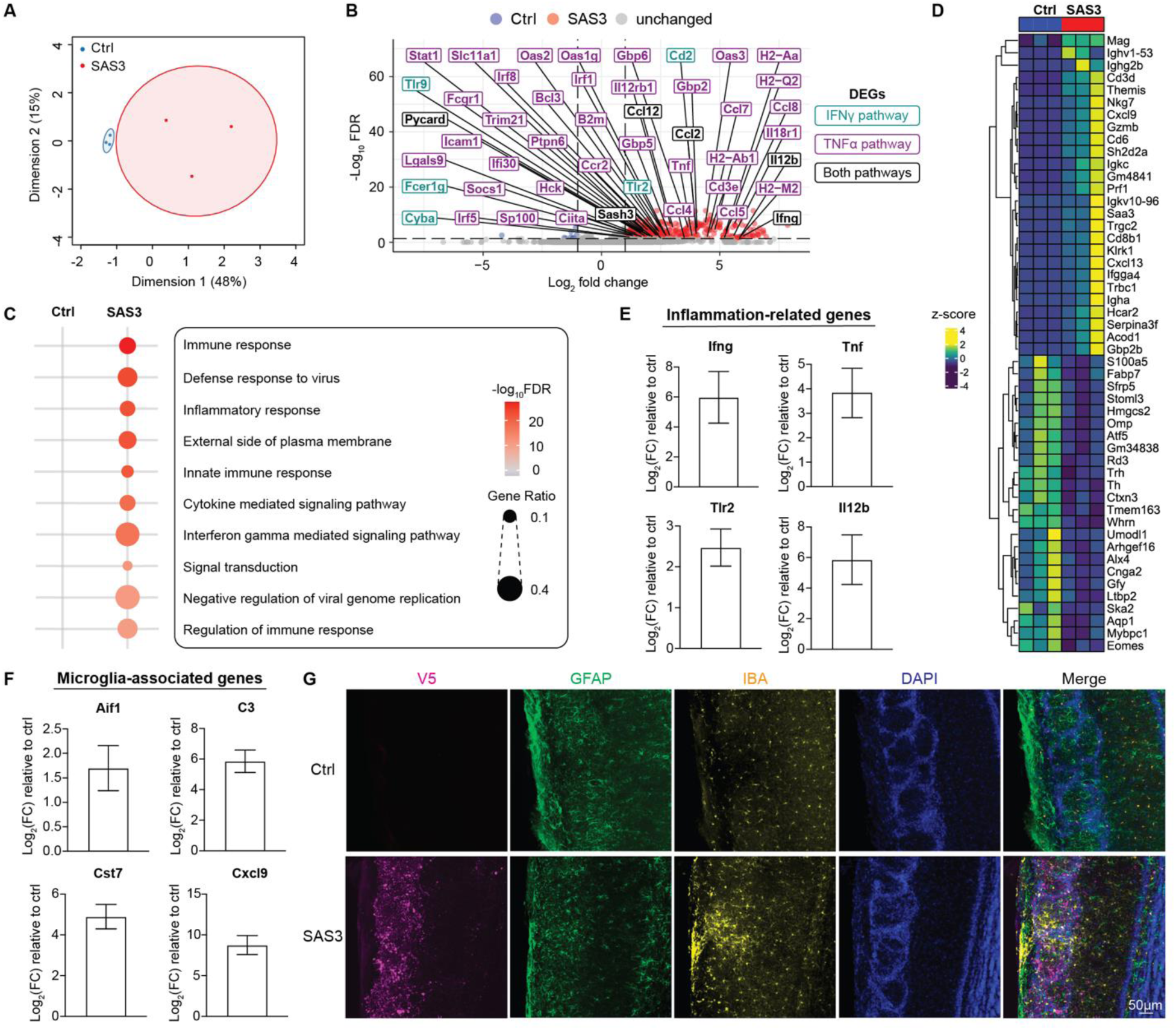
SAS activates microglia and triggers immune response in mice. **(A)** PCoA of transcriptomic data shows separation between SAS3-treated mice (transduced with hSyn-rtTA and TRE-SAS3 using AAV-PHP.eB) and controls (transduced with only hSyn-rtTA). n=3 biological replicates. **(B)** Volcano plot of DEGs with FDR < 0.05 and |log_2_ fold change (FC)| > 1. Genes belonging to the TNFα pathway are highlighted in magenta, IFN-γ pathway in cyan, and both in black. **(C)** Top 10 significantly altered GO pathways in SAS3-transduced mice indicate upregulation of pathways involved in inflammation. **(D)** Top 25 up- and downregulated genes. **(E)** Quantification of upregulation of representative genes of the inflammation-related IFN-γ, TNF-α, TLR, and interleukin pathways in SAS3-treated mice compared to controls. **(F)** Quantification of upregulation of representative genes involved in microglia activation. **(G)** Immunostaining of GFAP and IBA1 in the olfactory bulb, showing activated microglia clustered around SAS-induced aggregates. Scale bar equals 50 μm. Data are present as mean ± SEM.

We also observed a significant upregulation of genes associated with microglial activation in SAS-transduced mice (**Fig. 6F**). For example, IBA1, encoded by the *Aif1* gene, was markedly increased in SAS3-treated mice. CXCL9, a chemokine produced by microglia that plays a crucial role in neuroimmune responses, exhibited an approximately 500-fold upregulation. CST7, a cystatin that has been implicated in microglial activation under disease conditions and is linked to pathways involved in phagocytosis, cytokine/chemokine signaling, and antigen presentation, was also highly expressed. Additionally, complement protein C3, a key activator of microglia, showed a 60-fold increase. These changes strongly indicate a pronounced microglial activation state in SAS3-treated mice. Immunostaining of astrocytes (GFAP) and microglia (IBA1) confirmed that activated microglia clustered around SAS-induced aggregates, suggesting a targeted immune response to pathological protein accumulation (**Fig. 6G and S7B**). Collectively, these findings highlight the profound neuroinflammatory impact of SAS-induced aggregates, mirroring key aspects of PD pathology.

## Discussion

In this study, we present a synthetic biology platform that enables inducible, tunable, and cell-type-specific formation of intracellular aggregates of disease-relevant proteins in diverse model systems. This platform faithfully recreates hallmark features of human proteinopathies *in vitro* and *in vivo*, including inclusion localization, morphology and composition, transcriptional responses, and behavioral phenotypes. Using self-assembling α-synuclein (SAS), we established a PD model that recapitulates Lewy body-like pathology, dopaminergic neuron loss, aggregate propagation along the nigrostriatal axis, and progressive motor deficits. Importantly, SAS-induced aggregates incorporate mitochondria, lipids, ubiquitin, and p62, molecular signatures of bona fide Lewy bodies that have been difficult to reproduce in other disease models. The system also revealed prion-like seeding of endogenous αSyn and elicited robust glial responses, offering a powerful framework to study cell-nonautonomous aspects of disease progression. Moreover, by extending this approach to Tau and TDP-43, we demonstrated the platform’s generalizability across major neurodegenerative diseases. The observed differences in organelle interactions and toxicity of different protein aggregates illuminate potential mechanisms of selective vulnerability and disease divergence.

During our initial SAS construct screening, we discovered that different types of αSyn aggregates exhibit widely varying capacities to induce pS129-αSyn, a key disease marker. This raises critical questions about what structural and biochemical features define a toxic aggregate. Our findings suggest that local concentration, structural conformation, and context-dependent modifications play a decisive role. For example, SAS3-induced aggregates exhibit significantly higher toxicity than *SNCA* OE despite a lower total amount of αSyn (**Fig. 1C-D and S1B-C**). Similar observations have been made comparing αSyn PFFs and αSyn monomers. Structural differences appear to influence pathological outcomes, as different SAS constructs induce different levels of toxicity and phosphorylation (**Fig. S1A**). This aligns with studies showing that patient-derived αSyn PFFs exhibit different structural properties and varying neurotoxicity^44^. These observations imply that the structural conformation of αSyn aggregates may be a key determinant of disease progression and severity. Deciphering the structural determinants of aggregate toxicity may be essential for understanding disease progression and designing targeted interventions.

A major strength of this platform is its broad applicability across *in vitro* and *in vivo* experimental systems, including cell lines (**Fig. 1**), primary neurons (**Fig. 2**), hESC or iPSC-derived neurons, organoids (**Fig. 3&4**), and mice (**Fig. 5-6**). Notably, in hESC-derived neurons and brain organoids, SAS, SA-Tau and SA-TDP enable robust recapitulation of PD, AD, and ALS hallmarks, such as LB formation, extracellular Aβ production, and TDP-43 cytoplasmic inclusions, which previous models have failed to reproduce. This provides a compelling opportunity to integrate our self-assembling system with patient-derived iPSCs to dissect the impact of genetic variants and environmental factors on disease onset and progression. The system’s tunability also makes it well suited for high-throughput screens to identify disease modifiers and therapeutic compounds.

We provide direct evidence in primary neurons that SAS induces secondary nucleation of endogenous αSyn (**Fig. 4B&C**). This finding is further supported by the presence of V5-negative, pS129-positive aggregates in neurons in multiple, *in vitro* and *in vivo*, systems (**Fig. 2B; Fig. S2A; Fig. S3A; Fig. S4B; Fig. S6D**). Secondary nucleation is a critical driver of αSyn aggregation spread and amplification, yet its underlying mechanism remains unclear. Future studies are needed to dissect how primary nucleation triggers secondary nucleation and, more importantly, to identify strategies to halt secondary nucleation, which could be key to developing effective disease-modifying therapies for PD.

Neurodegenerative diseases often exhibit selective neuronal vulnerability^10^, and single-cell RNA sequencing (scRNA-seq) studies have identified distinct neuronal subtypes that are more susceptible to degeneration in PD^45,46^. In this study, we successfully demonstrated the ability to deliver SAS into specific neuron types non-invasively using AAV-PHP.eB. By targeting different neuronal subpopulations using cell-type-specific Cre lines or promoters, we can systematically investigate the roles of various neuronal populations in driving diverse neurodegenerative pathologies. This approach provides a powerful strategy to dissect the contributions of specific neuronal populations to disease progression, offering insights into why certain neurons are particularly susceptible and identifying potential therapeutic targets to protect these vulnerable populations.

While our primary focus was PD, αSyn aggregation is also central to other synucleinopathies, such as dementia with Lewy bodies (DLB) and multiple system atrophy (MSA)^9^. The SAS system can be readily extended to study these disorders, investigating potentially distinct or overlapping pathological mechanisms. More broadly, our synthetic approach provides a modular, programmable framework for modeling protein aggregation across a wide range of diseases. By applying this strategy to αSyn, Tau and TDP-43, we successfully recapitulated key features of PD, AD and ALS, respectively. By enabling side-by-side comparison of different disease-associated aggregates in a unified system, this platform opens the door to comparative pathology studies and mechanistic discovery across the neurodegenerative disease spectrum. In addition, protein aggregation is increasingly recognized in non-neurological diseases. For example, islet amyloid polypeptide (IAPP) aggregation cause β-cell dysfunction in type 2 diabetes, and aggregation of mutant p53 has been implicated in solid tumors. As such, this platform may hold value for a broad range of research communities beyond neuroscience.

Together, our work introduces a scalable, versatile and highly disease-relevant strategy to model and dissect protein aggregation. This “build-to-understand” framework sets the stage for fundamental discoveries into the mechanisms of aggregation-driven pathology and for the development of effective, disease-modifying therapies.

## Limitations of the study

While the self-assembling protein platform provides a powerful and programmable system to model intracellular protein aggregation and its pathological consequences, it has several limitations. First, although the aggregates recapitulate key disease hallmarks, the precise biophysical structures of the self-assembling aggregates may differ from endogenous protein aggregates found in patients. However, the heterogeneity of intracellularly forming aggregates, both in human patients and our system, present a technical challenge for high resolution structural characterization. In future studies, when technology and resources permit, in situ structural analysis of aggregates may offer more deeper insight into the relationship between aggregate structure and neurotoxicity. Second, our current study is intended as a conceptual and technical advance in the modeling of protein aggregation diseases. Due to the time and resource constrains, we selected SAS3 variant to conduct further studies based on its robust ability to recapitulate Parkinson’s disease pathology. However, this doesn’t preclude the possibility of other constructs, if tested at a longer duration or in different context, might yield aggregates that more closely resemble disease pathology. Finally, although we demonstrate *in vitro* applicability in human brain organoids and *in vivo* in the mouse, future studies are needed to assess the aggregates in more complex or long-lived models, such as non-human primates. Despite these limitations, our platform offers unprecedented spatiotemporal control and disease relevance, and we envision its broad utility in mechanistic investigations and therapeutic development.

## Resource availability

### Lead contact

Further information and requests for resources and reagents should be directed to, and will be fulfilled by, the lead contact, Dr. Viviana Gradinaru.

### Materials, data and code availability

The data, code, protocols, and key lab materials used and generated in this study are listed in a Key Resource Table alongside their persistent identifiers at Table S4. The bulk transcriptomics data generated and analyzed in this study have been deposited in the NCBI Sequence Read Archive (SRA) under BioProject accession number PRJNA1227536. The raw sequencing data and associated metadata are publicly available and can be accessed via the NCBI SRA database at https://www.ncbi.nlm.nih.gov/bioproject/PRJNA1227536. All code associated with analysis and plotting are available from https://github.com/jdthoang/FanY_2025_SAS.

## Acknowledgments

We thank Dr. Catherine Oikonomou for comprehensive editing of the manuscript, and Dr. Sarkis Mazmanian and Dr. Michael Elowitz for helpful discussions. We thank Patricia Anguiano for logistical assistance, and Karan Mahe and the rest of Gradinaru laboratory for critical feedback on the research and manuscript. This work was supported by the Office of Laboratory Animal Resources, CLOVER Center, and Millard and Muriel Jacobs Genetics and Genomics Laboratory at the California Institute of Technology. Figures contain images created with BioRender.com. Y.F. is supported by Caltech Chen Postdoc Innovator Grant and Parkinson’s Foundation Postdoctoral Fellowship award PF-PRF-1435603. This research was funded in part by Aligning Science Across Parkinson’s (ASAP-020495 to V.G.) through the Michael J. Fox Foundation for Parkinson’s Research (MJFF). For the purpose of open access, the authors have applied a CC BY public copyright license to all Author Accepted Manuscripts arising from this submission.

## Author contributions

Y.F. conceived the project, wrote the manuscript and prepared figures with input from all authors. V.G., G.M.C., C.S., J.D.H. assisted in manuscript preparation. Y.F. performed most experiments and collected and analyzed the data. J.D.H. performed the analysis of all RNA-seq data and contributed to animal experiments. C.S. performed all behavior assays, contributed to primary neuron, RNAseq and histology experiments, and assisted in many experiments. Y.B. contributed to cloning, RNAseq and organoid experiments. S.D. contributed to primary neuron and histology experiments. G.M.C. contributed to animal experiments. N.A. contributed to cloning and virus production. X.D. contributed to design of constructs. C.L. contributed to western blotting. Z.Q. contributed to virus production. E.D.M. contributed to primary neuron experiments. V.G. supervised all aspects of this study and funded the project.

## STAR Methods

### Key resources table (Table S4)

#### Animals

Animal husbandry and experimental procedures involving mice were done in accordance with protocols approved by the California Institute of Technology Institutional Animal Care and Use Committee (IACUC, protocol 1650) and the mice were handled in accordance with the principles and procedures of the Guide for the Care and Use of Laboratory Animals of the National Institutes of Health. Mice were group housed at 2-3 mice per cage with 13/11 light/dark cycles at ambient temperatures of 71–75°F and 30–70% humidity. C57BL/6J (Jackson Labs #026267), Snca^tm1.1Kluk/J^ (Jackson Labs #035412) and heterozygous B6.Cg-7630403G23Rik^Tg(Th-cre)1Tmd/J^ (Jackson Labs #008601) mice were used in experiments. Genotyping was performed by Transnetyx. AAVs were administered via retro-orbital injection during isoflurane anesthesia (1-3% in 95% O2/5% CO2, provided by nose cone at 1 L/min), at 1 × 10^12^ viral genomes (v.g.) per animal. A detailed protocol for systemic AAV administration through retro-orbital injection is available on protocols.io (dx.doi.org/10.17504/protocols.io.36wgqnw73gk5/v1). Mice were randomly assigned to different conditions.

#### Cell lines

HEK 293 cells (ATCC, CRL-3216) were used in SAS screening and characterization. The human embryonic stem cell line (hESC) line CSES07 (Cedars Sinai, CVCL_B818) was used to derive both monolayer neurons and organoids. CSES07 was obtained from Cedars Sinai Medical Center with a verified normal karyotype and was contamination-free. All cell lines were authorized for use under the supervision of the California Institute of Technology Institutional Biosafety Committee (IBC #22-182).

#### SAS library, SA-Tau and SA-TDP design

To investigate whether intracellular αSyn aggregation can recapitulate Parkinsonian pathology, we designed a library of self-assembling synuclein (SAS) constructs. Each construct consisted of wild-type (WT) αSyn (UniProt ID: P37840-1) fused to selected self-assembling proteins (SAPs). SAPs were chosen based on structural evidence indicating terminal accessibility and the nature of their assemblies, as revealed by structural studies. 1POK(E239Y)^24^, DHF40, and DHF46^22^ were selected based on known filament formation (PDB entries 5LP3 and 6E9R), while 2VYC (K491L, D494L, D497L)^47^ and gamma-prefoldin (γPFD)^25^ were selected for their punctate assembly characteristics (PDB entry 6VY1). Additionally, cytoskeletal proteins β-tubulin (PDB: 3J2U) and β-actin (PDB: 6DJO) were included to evaluate intracellular fiber-mediated effects. Several constructs also integrated mitochondrial targeting proteins (TOMM5, TOMM20) to explore how spatial localization influences aggregation outcomes. All fusion junctions incorporated flexible glycine-rich linkers (GGSGGTGG) and V5 epitope tags for enhanced detection and characterization. SA-Tau and SA-TDP construct consisted of human Tau (isoform 2N4R) or TDP-43 fused with 1POK(E239Y). Detailed information on SAS, SA-Tau and SA-TDP constructs is listed in Supplemental Table 1.

#### SAS screening in HEK cells

SAS plasmids were transfected into HEK cells using Lipofectamine 3000 (ThermoFisher, L3000008) following the manufacturer’s instructions. Briefly, 100K HEK cells were plated in 24-well plates. The next day, 0.4 μg SAS plasmids and 0.1 μg rtTA plasmids were transfected into each well. Cells were incubated with lipofectamine overnight. The medium was switched to 5% FBS in DMEM/F12 the next morning, and 10 μg/mL of doxycycline (Sigma, Cat # D9891-10G) was added into the medium. After 3 days of doxycycline treatment, cells were fixed and stained. For protocol, see dx.doi.org**<u>/10.17504/protocols.io.6qpvr97wzvmk/v1</u>**

#### Immunostaining of cultured cells

At predetermined time points, cells were removed from the incubator, washed once with phosphate-buffered saline (PBS), and fixed with 4% paraformaldehyde for 10 minutes at room temperature (RT). After washing 3 times with PBS, the blocking buffer containing 1xPBS, 0.03% Triton X-100 (PBST), and 5% normal donkey serum was applied and incubated at RT for 60 minutes. Primary antibodies were diluted in fresh blocking buffer, added on the cell, and incubated overnight at 4°C. The next day, the antibody solution was washed off with PBST for 10 minutes, 3 times. Secondary antibodies were applied for 60 minutes and then washed off with PBST for 10 minutes, 3 times. After the final wash, cells remain in PBST for imaging. Antibodies and reagents used were listed in the key resource table. For protocol, see dx.doi.org**<u>/10.17504/protocols.io.4r3l29eb4v1y/v1</u>**

#### Western blotting

Western blotting was performed using the iBind Automated Western System following the manufacturer’s instructions. Briefly, cells were collected with cold DPBS and lysed with RIPA buffer. Total protein concentration was measured using Qubit Protein BR Assay kit and Qubit4. An equal amount of protein was run in Bio-Rad 4-20% TGX stain-free gels and then transferred to a PVDF membrane. Membranes were fixed with 4% paraformaldehyde for 10 minutes before blotting. Protein immunodetection was performed with indicated antibodies using the iBind Automated Western System. Signals were detected with Max Western ECL and imaged with a ChemiDoc imaging system. Antibody information and reagents used are listed in the key resource table (**Table S4**). For protocol, see dx.doi.org**<u>/10.17504/protocols.io.j8nlk96p6v5r/v2</u>**

#### Viral vector production

AAV packaging and purification were carried out as previously described^48^. The viruses for *in vitro* experiments were produced by suspension HEK cells. In brief, recombinant AAV was produced by triple transfection of cells in suspension using the VirusGEN AAV Kit with RevIT enhancer (Mirus Bio, MIR 8007) at a molar ratio of transgene: AAV capsid: pHelper = 1:2:0.5, as per the manufacturer’s instructions. The total DNA amount was 2 µg per mL of cells. Virus-producing cells and medium were harvested 72 hours post-transfection. The viruses used in animal experiments were produced by adherent HEK cells. In brief, HEK cells are transiently transfected using polyethyleneimine (PEI) at a molar ratio of transgene: AAV capsid: pHelper = 1:4:2. 120 hours after transfection, the producer cells were harvested, and cell culture media was collected at 72- and 120-hours post-transfection. Following cell lysis and nuclease digestion, viruses from suspension cells and adherent cells were purified using iodixanol step gradient columns followed by ultracentrifugation, as previously detailed^48^. The purified AAVs were quantified through droplet digital PCR (ddPCR) following Addgene’s protocol with modifications^49^. Briefly, AAV samples were first serially diluted to achieve a suitable concentration for ddPCR analysis. The PCR reaction mixture was prepared using ddPCR EvaGreen Supermix (Bio-Rad), primers for WPRE or ITRs, the diluted AAV sample, and nuclease-free water. The mixture was then partitioned into droplets using the QX200 Droplet Generator, and the droplets were transferred to a 96-well plate for PCR in a thermocycler. After cycling, the droplets were read using a QX200 Droplet Reader, and the concentration of the viral genome was analyzed using QuantaSoft software. The viral genome titer in original sample was then back calculated using the dilution factors. The final viral preparation was stored in DPBS with Pluronic F-68 to prevent adhesion loss. The reagents used are listed in the key resource table (**Table S4**). For protocol, see dx.doi.org**<u>/10.17504/protocols.io.ewov1dpeovr2/v1</u>**

#### Primary neuron preparation

The day before the procedure, 10× HBSS (Thermo Fisher, Cat # 14065056) was diluted to 1× using sterile distilled water (Invitrogen, Cat # 10977015) and pre-cooled to 4°C. Glass inserts (Neuvitro Corporation, Cat # GG-12-15 Pre) were placed in 24-well plates and were coated with Poly-D-lysine (Millipore Sigma, Cat # A-003-E) overnight. Plating media was prepared using the BrainPhys Neuronal Medium SM1 Kit (Stem Cell Technologies, Cat # 05792), 1× Glutamax, (Thermo Fisher,Cat # 35050061), 1× Penicillin-Streptomycin (Thermo Fisher, Cat # 15140122,), and 1× l-glutamine (Bio-Techne, Cat # 0218). Pregnant mice from JAX were euthanized on day 17 of pregnancy. Embryos were removed, decapitated, and brains were isolated in cold HBSS. Cortices and hippocampi were dissected, and hippocampi were quartered and placed into 15 mL tubes. Papain (Sigma Aldrich, Cat # P3125-250MG) was added for digestion, followed by washing with a stopping medium containing HBSS 1× and 10% Fetal Bovine Serum. Hippocampi were dissociated by gentle pipetting, and supernatants with dissociated neurons were collected. After centrifugation, cells were resuspended in plating media, counted, and seeded at 200,000 cells/well on Poly-D-lysine-coated inserts. After 72 hours, the media was replaced with a combination of 500 mL Neurobasal Medium (Thermo Fisher, Cat # 21103049), 2 mL B27 Supplement (Thermo Fisher, Cat # 17504044), and 0.5 mM L-glutamine (Bio-Techne, Cat # 0218). Virus was then added for indicated times and doses. For protocol, see http://dx.doi.org/10.17504/protocols.io.5qpvo98ozv4o/v1.

#### Cortical neurons and organoid differentiation

hESCs were plated on 6-well plates coated with Vitronectin (Gibco, Cat # A14700) at 1:100 diluted in DPBS (Gibco, Cat #14190250) and maintained in E8 medium (Thermo Fisher, Cat #A1517001). The E8 medium was changed every day, and the cells were passaged every 3-5 days at 70-85% confluence. The cells were passaged using EDTA dissociation as described previously^50^. To induce differentiation, hESCs were dissociated at 70%-80% confluence using EDTA dissociation buffer and replated as a single cell suspension on Matrigel-coated 24-well plates at a density of 100K cells/cm^2^ in E6 medium (Thermo Fisher, Cat #A1516401) containing 10 μM ROCK-inhibitor (Y-27632; R&D, Cat #1254). The cells were kept in E6 medium + Y-27632 overnight. The next day, the medium was switched to a neuroectoderm-inducing medium. The cells were kept in E6 medium containing 10 μM SB431542 (R&D, Cat #1614), 100 nM LDN139189 (R&D, Cat #6053), and 500 nM XAV (R&D, 3748) for the first 4 days and then switched to E6 medium containing 10 μM SB and 100 nM LDN for 6 days. Then the cells were switched to progenitor expanding medium, containing neurobasal (NB) medium supplemented with l-glutamine (Gibco, Cat #25030-164), N2 (Stem Cell Technologies, 07156), B27 (Life Technologies, Cat #17504044) and NEAA (Sigma, Cat #M7145) for 10 days. At D20, the neuron progenitors were dissociated into single cells and replated in a V-bottom ultra-low attachment 96-well plate at 100K per well to form neuron spheres. After 4 days, the spheres were transformed to low attachment 10 cm dishes and placed on a shaker for long-term culture and maturation in cortical neuron medium containing NB/N2/B27/Glu/NEAA, ascorbic acid (200 μM, Sigma, 4034-100g), dbcAMP (500 μM, Sigma, Cat #D0627), BDNF (20 ng/mL, R&D, Cat #248-BDB) and GDNF (20 ng/mL, Peprotech, Cat #450-10)^51^. Organoids were cultured in suspension for 90 days before adding viruses. For protocol, see dx.doi.org/10.17504/protocols.io.3byl4wrzovo5/v1

#### Open field test (OFT)

Mice were tested in the OFT bi-weekly. The open-field apparatus consisted of four square arenas (27 cm × 27 cm), with a camera (EverFocus, EQ700) placed 1.83 m above the floor of the arenas. EthoVision XT was used to capture and subsequently analyze animal locomotion. Each trial consisted of a 2 min habituation period, followed by a 10 min test period. To avoid confounds due to odors from non-cagemates, only animals from the same cage were recorded simultaneously. Behavior equipment was disinfected and deodorized between each animal. The distance traveled over the course of the experimental period was determined. For protocol, see dx.doi.org**<u>/10.17504/protocols.io.5qpvo972xv4o/v1</u>**

#### Beam crossing assay

To measure skilled locomotion using the narrowing beam assay, a clear plexiglass beam consisting of three 25 cm segments (widths 3.5 cm, 2.5 cm, and 1.5 cm) was elevated above the table surface using empty clean cages. At the narrow end, an empty cage was placed on its side, and bedding from the animal’s home cage was placed inside. A white light was also placed over the broad end to motivate animals to move across the beam. For each trial, animals were placed at the end of the widest segment, with all 4 limbs touching the beam surface. Each trial was recorded with a video camera placed to the side and perpendicular to the beam’s length, affording a view of both left and right hindlimbs. A trial was considered complete once the animal had traversed the beam, without turning around, and entered the goal cage. Once an animal had completed three trials, the session was completed. Beam crossing was analyzed using BORIS software, measuring the time taken to cross the beam and the number of foot slips. For protocol, see dx.doi.org**<u>/10.17504/protocols.io.n92ldr2jxg5b/v1</u>**

##### Tissue harvesting and processing

Animals were transcardially perfused with 30 mL of ice-cold heparinized 1× PBS, and brains were dissected. One hemisphere of the brain was used for RNAseq. For analysis of fluorescent protein expression, one hemisphere of the brain was submerged in ice-cold 4% PFA formulated in 1× PBS and fixed for 24-48 hr at 4°C. Brains were subsequently cryoprotected at 4°C in a solution containing 30% (w/v) sucrose for 72 hr. Brains were flash-frozen in O.C.T. Compound (Scigen, Cat # 4586) using a dry ice-ethanol bath and kept at -80°C until sectioning. For protocol, see dx.doi.org**<u>/10.17504/protocols.io.ewov1d5xovr2/v1</u>**.

#### Immunostaining of mouse tissue

Brains were sliced at 100 μm using a cryostat (Leica Biosystems, CM1950) and collected in 1× PBS. Sections were stored in PBS supplemented with 0.02% Azide. For protocol, see dx.doi.org**<u>/10.17504/protocols.io.x54v9rkzpv3e/v1</u>**.

For immunostaining, brain sections were blocked with 5% normal donkey serum in 0.1 % PBST for 90 mins, then incubated with primary antibody solution prepared in 0.1 % PBST supplemented with 5% normal donkey serum overnight on a shaker at 4°C. After washing in 0.1 % PBST (3 × 30 min), the sections were incubated with secondary antibodies overnight on a shaker at 4°C and then washed 3 x 30 min in 0.1 % PBST. The tissues were mounted on glass slides with Prolong Diamond Antifade mounting media (Thermo Fisher Scientific, Cat # P36970). Imaging was performed with a spinning disk confocal microscope. Antibodies are listed in the Key Resource Table (Table S4). For protocol, see https://dx.doi.org/10.17504/protocols.io.kqdg3q7ypv25/v1

#### RNA extraction

RNA extraction from cells was performed using the Direct-zol RNA Miniprep kit (Zymo, Cat # R2051). Briefly, the cells were removed from incubator, washed once with DPBS, then 350 µL of TRIzol was added directly to each well to collect the cells. Subsequent RNA extraction was done following the manufacturer’s instructions. For protocol, see dx.doi.org**<u>/10.17504/protocols.io.6qpvr9r2zvmk/v1</u>**

RNA extraction from organoids was performed using the QIAshredder kit (Qiagen, Cat # 79654) and miRNeasy Mini Kit (Qiagen Cat # 217004). Briefly, 3 organoids were harvested for each RNA extraction and 350 µL of Buffer RLT from RNeasy Mini Kit + 3.5 µL β-mercaptoethanol were added to lyse the organoid. Subsequent RNA extraction was done following the manufacturer’s instructions. For protocol, see dx.doi.org**<u>/10.17504/protocols.io.81wgbr95qlpk/v1</u>**

RNA extraction from mouse brain tissue was performed using the miRNeasy Mini Kit (Qiagen catalog, Cat # 217004) on a QIAcube Connect (Qiagen). After perfusion with cold PBS, the entire olfactory bulb was dissected and placed in TRIzol directly and processed immediately for RNA extraction following manufacturer’s instructions. For protocol, see dx.doi.org**<u>/10.17504/protocols.io.eq2ly618pgx9/v1</u>**.

#### RNA sequencing (RNA-seq)

Bulk RNAseq was performed by the Millard and Muriel Jacobs Genetics and Genomics Laboratory at the California Institute of Technology. RNA integrity was assessed using the RNA 6000 Pico Kit for Bioanalyzer (Agilent Technologies, Cat # 5067-1513), and mRNA was isolated from ∼1 μg of total RNA using the NEBNext Poly(A) mRNA Magnetic Isolation Module (NEB, Cat #E7490). RNA-seq libraries were constructed using the NEBNext Ultra II RNA Library Prep Kit for Illumina (NEB, Cat. No. / ID: E7770) following the manufacturer’s protocols. Briefly, mRNA was fragmented to an average size of 200 nt by incubating at 94°C for 15 min in the first strand buffer. cDNA was then synthesized using random primers and ProtoScript II Reverse Transcriptase followed by second strand synthesis using NEB Second Strand Synthesis Enzyme Mix. The resulting DNA fragments were end-repaired, dA-tailed and ligated to NEBNext hairpin adaptors (NEB, Cat. No. / ID: E7335). Following ligation, adaptors were converted to the “Y” shape by treating with USER enzyme, and DNA fragments were size selected using Agencourt AMPure XP beads (Beckman Coulter # A63880) to generate fragment sizes between 250-350 bp. Adaptor-ligated DNA was PCR amplified followed by AMPure XP bead clean up. Libraries were quantified using a Qubit dsDNA HS Kit (ThermoFisher Scientific, Cat. No. / ID: Q32854), and the size distribution was confirmed using a High Sensitivity DNA Kit for Bioanalyzer (Agilent Technologies, Cat. No. / ID: 5067-4626). Libraries were sequenced on an Illumina NextSeq2000 in paired end mode with a read length of 50 nt and sequencing depth of 25 million reads per library. Base calls and FASTQ generation were performed with DRAGEN 4.2.7.

#### Transcriptomics data analysis

Raw sequencing reads were processed and aligned by Rsubread package v2.16.1 (https://bioconductor.org/packages/release/bioc/html/Rsubread.html, RRID:SCR_016945) in R (v4.3.2) to align trimmed reads. For human cortical organoids, reads were aligned to the GRCh38.p14 reference genome, while mouse brain tissues were aligned to the GRCm39 reference genome. The corresponding gene annotation files (GTF) were used with featureCounts (https://subread.sourceforge.net/featureCounts.html, RRID:SCR_012919) to generate gene level counts for each dataset. Data were subsequently normalized with DESeq2 v1.42.1 (https://bioconductor.org/packages/release/bioc/html/DESeq2.html, RRID:SCR_015687) and principal coordinate analysis was subsequently performed by multidimensional scaling to assess sample level clustering using the limma package (v3.60.4, RRID:SCR_010943). DESeq2 was then used to assess differential gene expression between rtTA and SAS3 samples. A two-tailed false discovery rate (FDR) < 0.05 (original Benjamini-Hochberg method) determined statistical significance and |log_2_(FC)| > 1.0 was selected to identify differentially expressed genes (DEGs). Pathway analysis by statistical overrepresentation was performed with DEGs separately for upregulated and downregulated genes with the Gene Ontology (GO) database using the Rapid Integration of Term Annotation and Network (RITAN) package v1.26.0577 (https://bioconductor.org/packages/release/bioc/html/RITAN.html). A two-sided adjusted p-value (q-value) threshold of 0.05 (Benjamini-Hochberg method) was used to determine statistically overrepresented pathways.

#### Image analysis

Images were processed and analyzed with imageJ2 V2.14.0/1.54f. The “*Analyze Particles*” and “*Nucleus Counter*” plugins were used for aggregate size and number analysis; “*Analyze Skeleton*” was used for primary neuron neurite length quantification. The fluorescence intensity measurement was used to determine striatal TH projection intensity.

#### Quantification and statistical analysis

Data are presented as mean ± SEM and were derived from at least three independent experiments. Information on replicates (n) is given in figure legends. Statistical analysis was performed using the unpaired t-test, also known as Student’s t-test (comparing two groups) or ANOVA with Dunnett test (comparing multiple groups against control). Distributions of raw data approximated normal distributions.

---

**Fig. S1.**
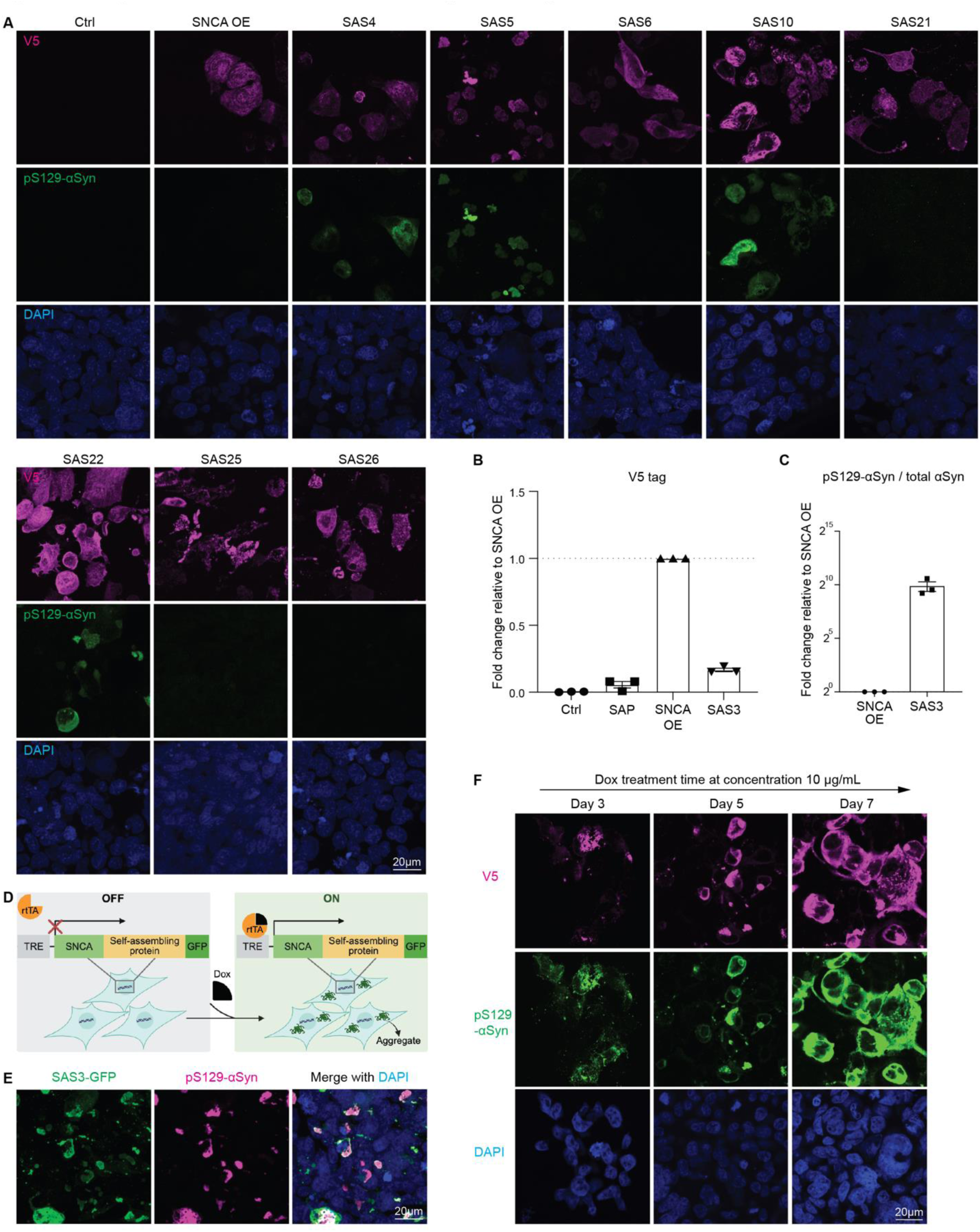
Screening of SAS constructs in HEK cells and tunability of the SAS system. **(A)** Screening in HEK cells to identify SAS constructs that form αSyn aggregates and induce substantial Ser129 phosphorylation of αSyn. **(B-C)** Quantification of western blot for total expression (V5 tag) and pS129-αSyn show that although SAS3 expression is lower than *SNCA* overexpression (*SNCA* OE), it induces 1000-fold greater phosphorylation, indicating that the enhanced pS129-αSyn signal is due to aggregation rather than protein level. n=3 biological replicates. **(D)** Design of SAS construct with GFP reporter. **(E)** Immunostaining of pS129-αSyn with GFP reporter after 3 days of dox induction, demonstrating that the GFP tag does not alter SAS behavior. **(F)** Immunostaining of V5 tag and pS129-αSyn after 3, 5, and 7 days of dox-induced SAS3 expression, demonstrating that aggregate size and level of pS129-aSyn increase with induction time. Scale bars equal 20 μm. Data are presented as mean ± SEM.

**Fig. S2.**
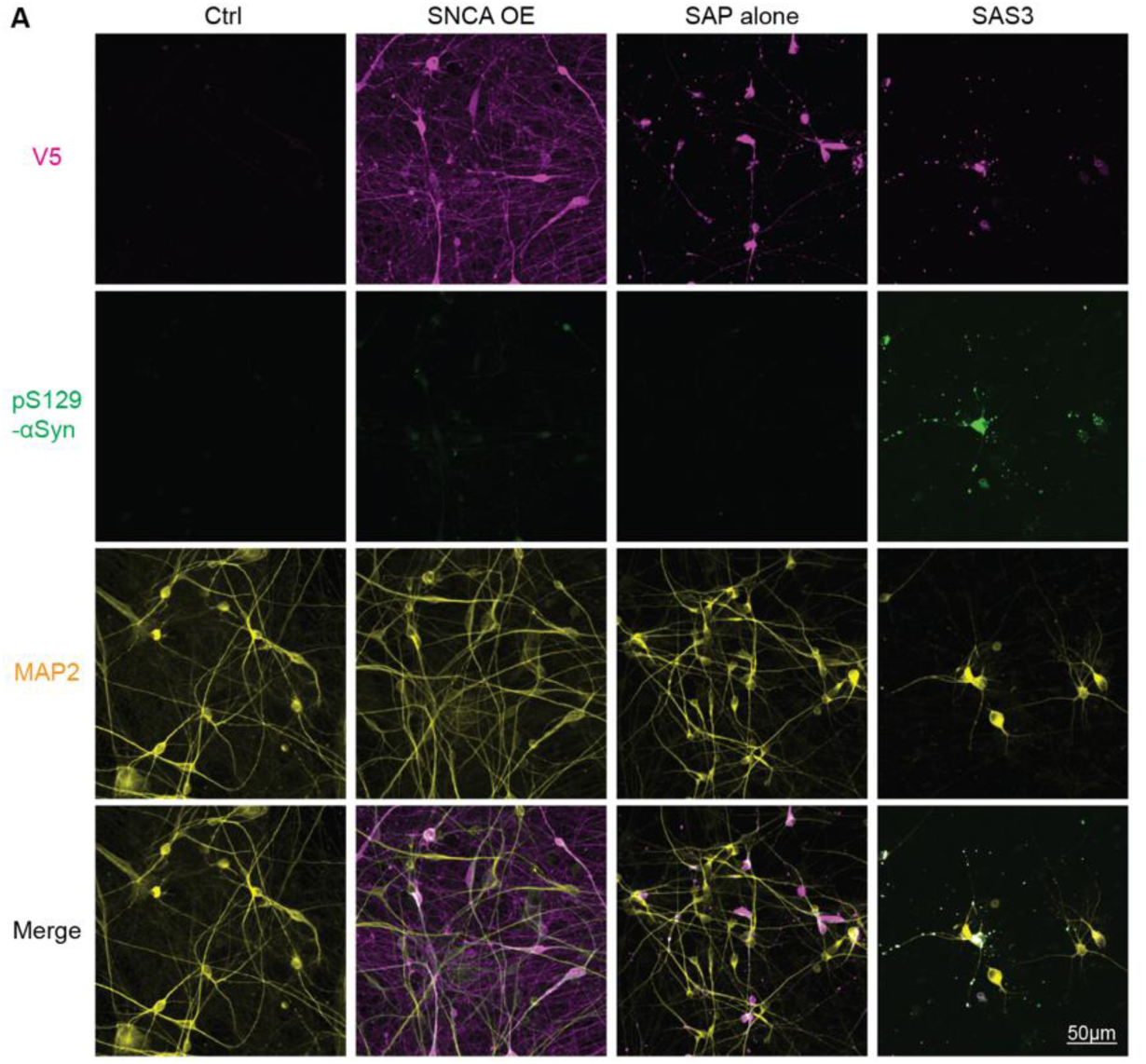
SAS induces αSyn aggregation and neurite retraction in hESC-derived neurons. **(A)** Immunostaining of V5 tag and pS129-αSyn in transduced hESC-derived human cortical neurons after 3 days of dox treatment shows that SAS3 induces αSyn aggregation, strong αSyn phosphorylation, and neurite retraction. Scale bar equals 50 μm.

**Fig. S3.**
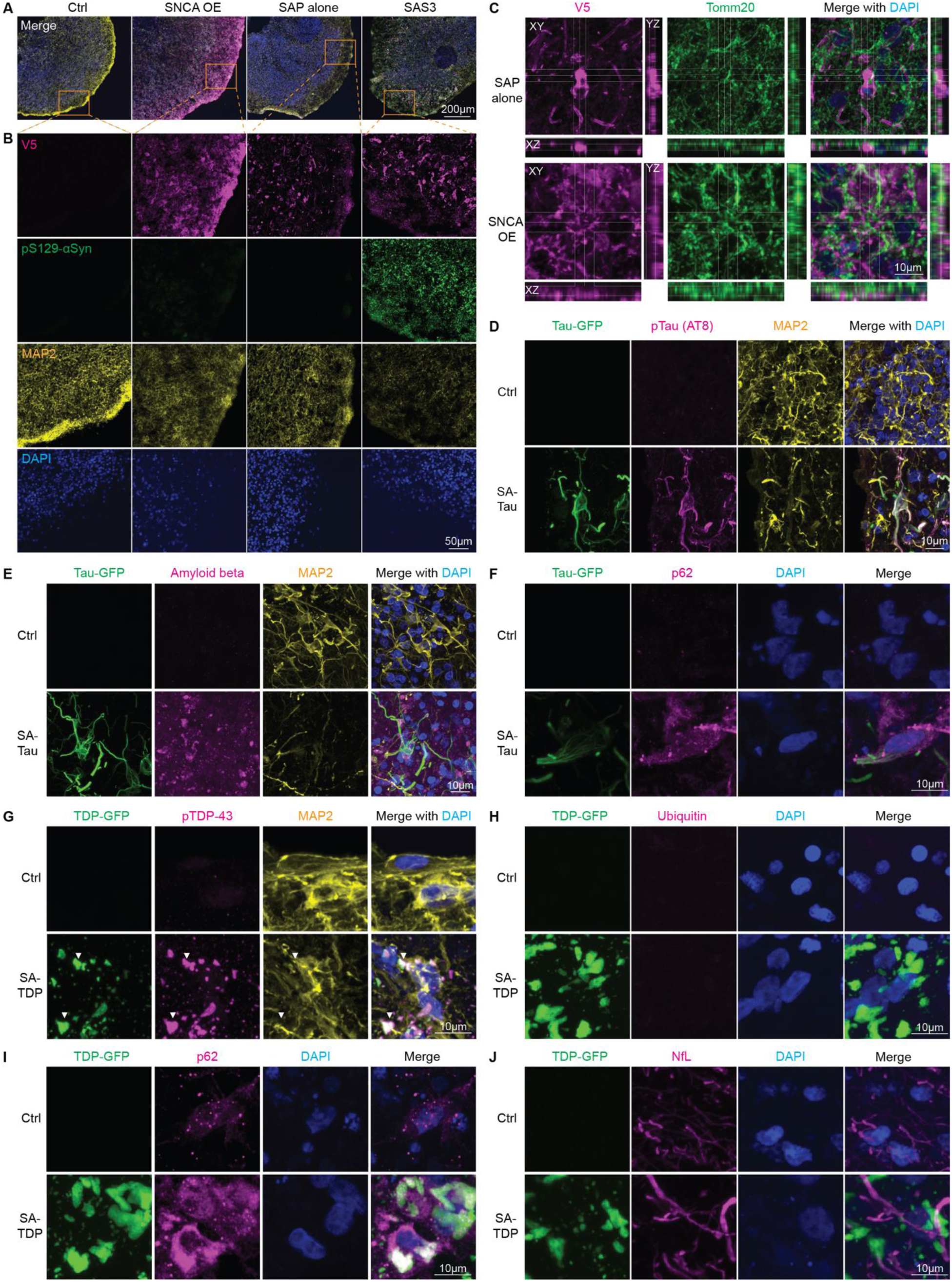
Self-assembling system reveals distinct organelle interactions and pathological signatures of αSyn, Tau, and TDP-43 aggregates. (A-B) Immunostaining of V5 tag and pS129-αSyn in transduced human cortical organoids after 3 days of dox treatment. Compared to *SNCA* OE and SAP alone, only SAS3 induces αSyn aggregation, intense αSyn S129 phosphorylation, and neuron degeneration. Scale bar equals 200 μm and 50 μm in enlargement. **(C)** Z-stack scanning of organoids stained for V5 tag and the mitochondrial marker TOMM20, showing that *SNCA* OE and SAP construct-produced proteins do not colocalize with mitochondria, shown by XY (main), XZ (bottom panel) and YZ (right panel) projections. **(D)** Immunostaining of GFP, pTau (AT8 antibody), and MAP2 in human brain organoids shows robust Tau aggregation and pTau, as well as severe neuron degeneration. **(E)** Immunostaining for amyloid beta in human brain organoids expressing SA-Tau reveals Tau tangles promote the production of extracellular amyloid beta. **(F)** SA-Tau expressing cells exhibited elevated accumulation of p62 signal. **(G)** Representative images of human brain organoids expressing SA-TDP show cytoplasmic TDP-43 inclusions with strong pTDP-43 (arrowheads). Compared with SAS and SA-Tau, human brain organoids expressing SA-TDP only shows mild neurodegeneration indicated by MAP2 staining. **(H)** Ubiquitin staining shows SA-TDP-induced aggregates are not ubiquitinated, unlike αSyn or Tau aggregates. **(I)** Accumulation of p62 in SA-TDP expressing cells indicates elevated autophagy. **(J)** SA-TDP-induced aggregates do not colocalize with neurofilament light chain (NfL). Scale bar equals 10 μm in **B-J**.

**Fig. S4.**
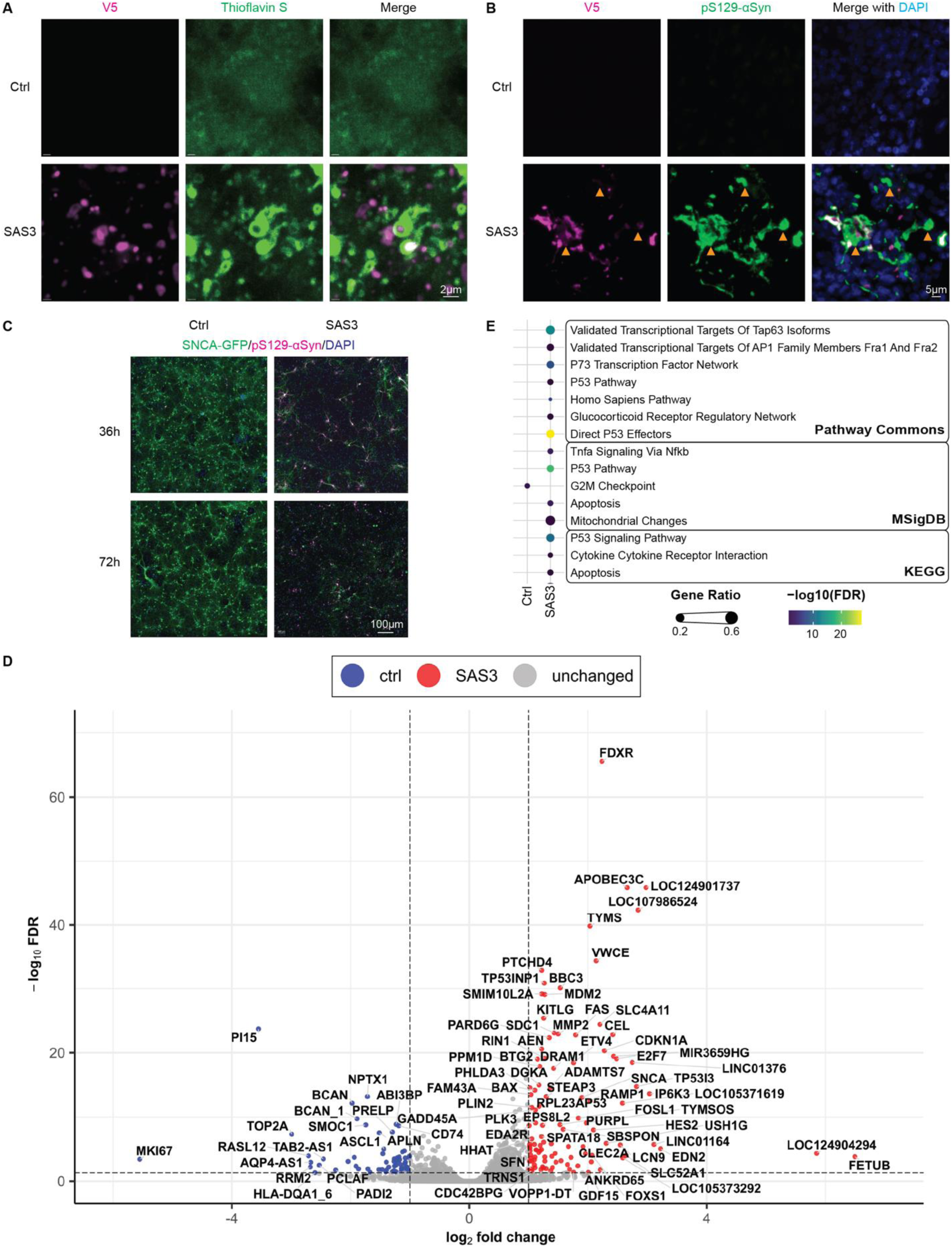
SAS forms LB-like inclusions and nucleates aggregation of endogenous αSyn, triggering neuronal inflammation and death. **(A)** Organoids stained with thioflavin S demonstrate the presence of β-sheet structures within and around SAS-induced aggregates. Scale bar equals 2 μm. **(B)** Organoids stained for V5 tag and pS129-αSyn reveals the presence of V5-negative and pS129-αSyn-positive aggregates (arrowheads), indicating the formation of endogenous αSyn aggregates. Scale bar equals 5 μm. **(C)** Immunostaining after 36 and 72 hours of dox treatment indicates that neurons expressing SAS3 display degeneration at 36 hours which further worsens by 72 hours. Scale bar equals 100 μm. **(D)** Volcano plot of transcriptomic data labeling all DEGs with FDR < 0.05 and |log2 fold change (FC)| >1. Genes upregulated in SAS-treated organoids are shown in red and those downregulated in blue. **(E)** Pathway analysis of DEGs using different databases reveals that SAS3-treated organoids are characterized by significant up-regulation of pathways associated with apoptosis, inflammation, and mitochondrial changes.

**Fig. S5.**
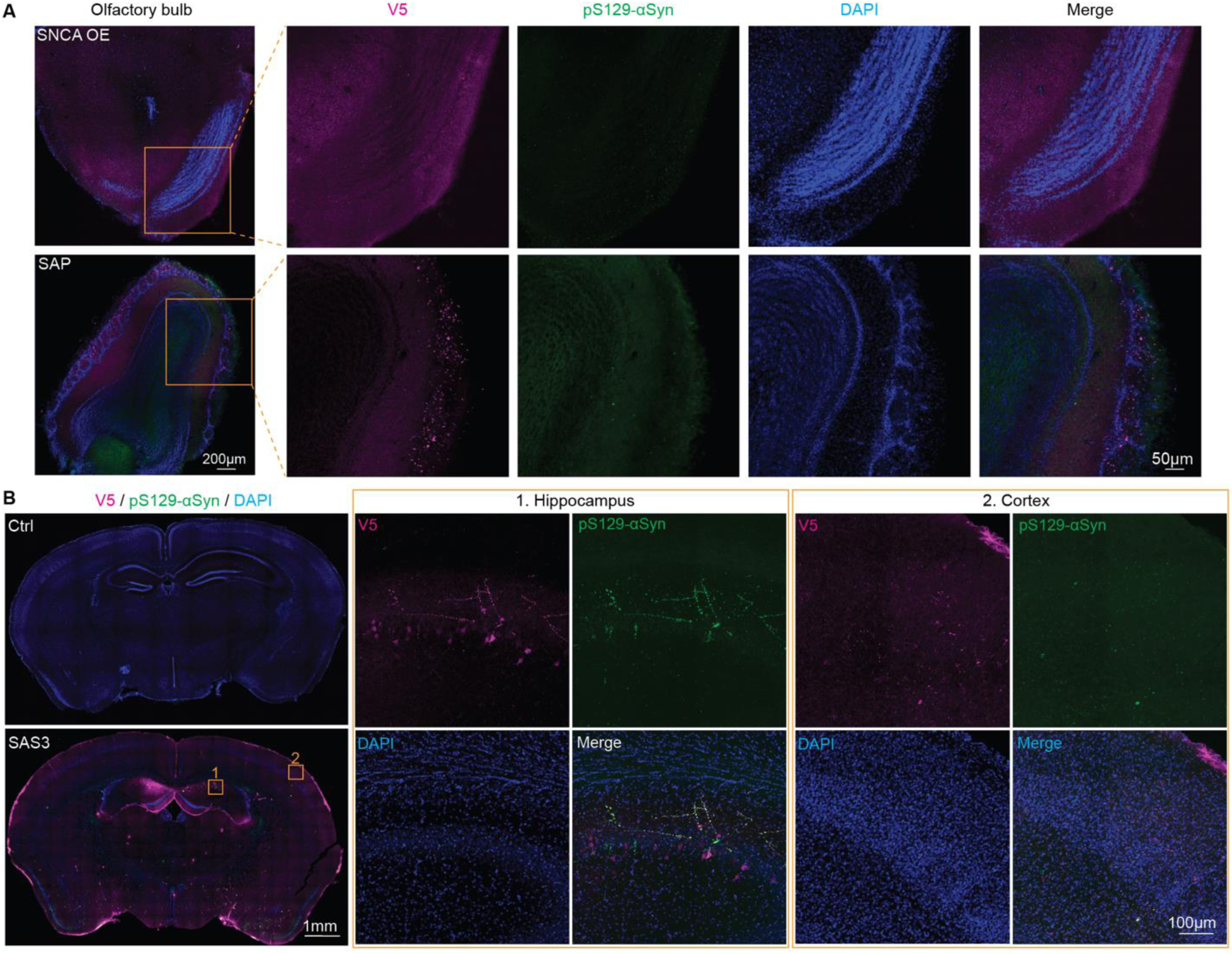
Systemic delivery of SAS induces PD-relevant histopathology. **(A)** Immunostaining for the V5 tag and pS129-αSyn in the olfactory bulb of C57BL/6J mice, showing that *SNCA* OE and SAP-alone constructs do not induce strong αSyn phosphorylation in olfactory bulb. Scale bar equals 200 μm and 50 μm in enlargement. **(B)** Immunostaining for the V5 tag and pS129-αSyn in the hippocampus and cortex of SAS3-treated or control mice, showing aggregates and strong αSyn phosphorylation in neurons induced by SAS3. Scale bar equals 1 mm and 100 μm in enlargement.

**Fig. S6.**
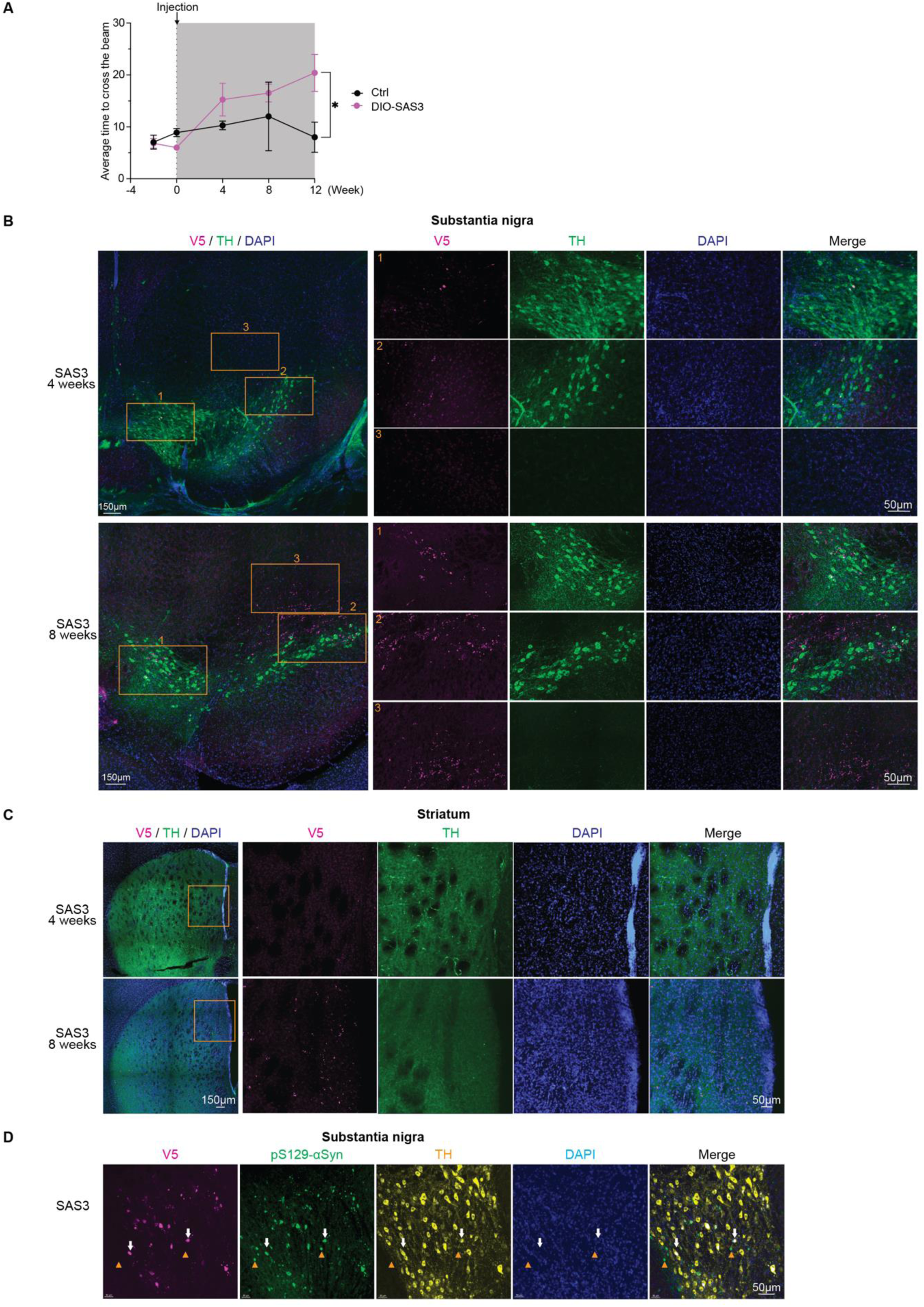
Dopaminergic neuron-targeted SAS recapitulates PD progression. **(A)** After 4 weeks of dox treatment, DA neuron-targeted SAS3 mice exhibited significant motor impairments in the beam crossing assay, requiring more time to cross the beam. **(B)** Immunostaining for the V5 tag and TH in the SN region of SAS3-treated TH-Cre mice at 4 weeks and 8 weeks, with enlarged images of the ventral tegmental area (VTA, area 1), SNpc (area 2), and non-TH+ region (area 3). Scale bars equal 150 μm and 50 μm in enlargements. **(C)** Immunostaining for the V5 tag and TH in the striatum of SAS3-treated TH-Cre mice at 4 weeks and 8 weeks, indicating progressive aggregate accumulation and potential spreading along the nigrostriatal pathway. Scale bar equals 150 μm and 50 μm in enlargement. **(D)** Immunostaining for V5, pS129-αSyn, and TH in the SNpc reveals that not only SAS3-expressed proteins (arrows) but also endogenous αsyn (arrowheads) formed aggregates and became phosphorylated. Scale bar equals 50 μm. Data are presented as mean ± SEM; statistical analysis was performed using unpaired t test; *p<0.05.

**Fig. S7.**
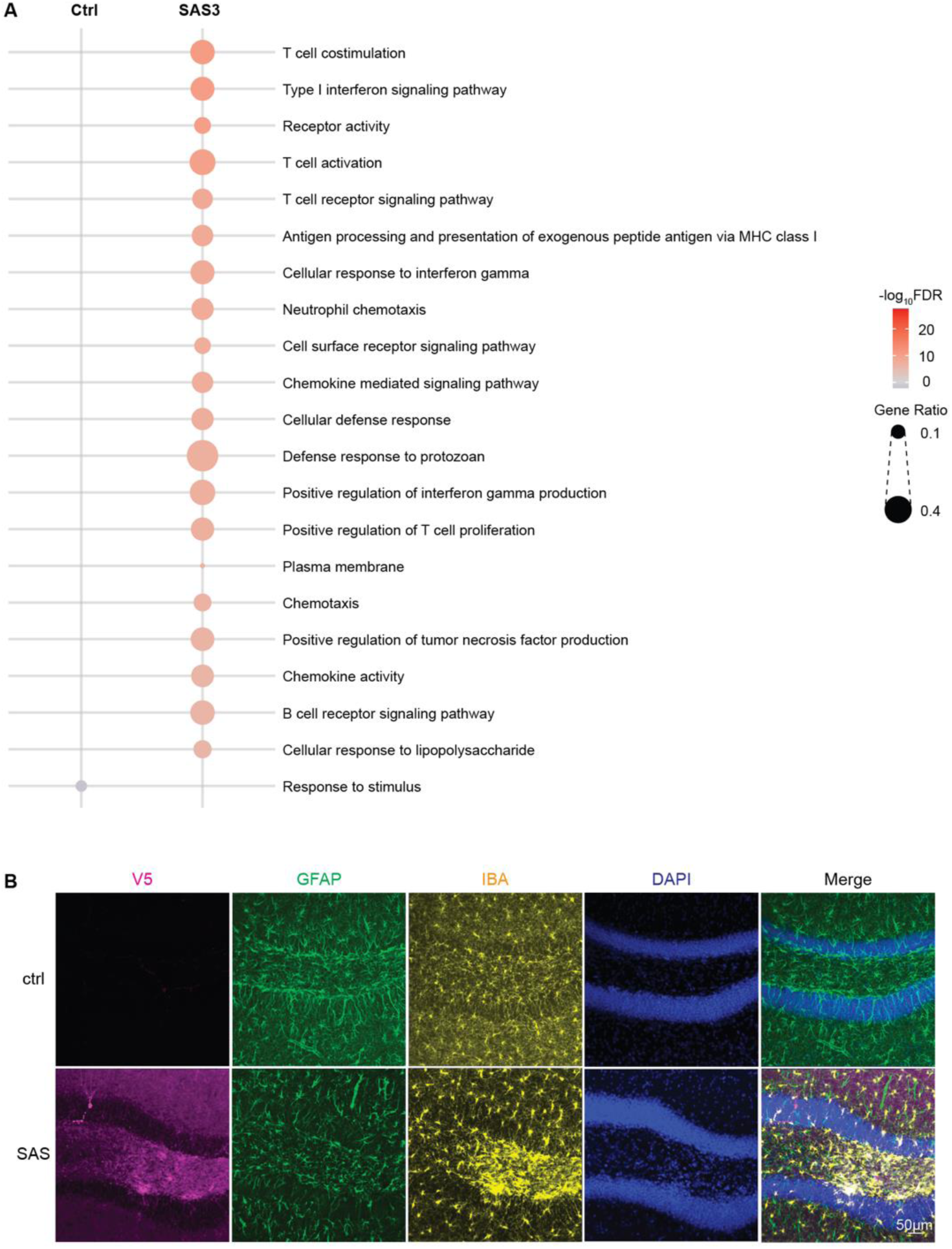
SAS activates microglia and triggers immune response in mice. **(A)** Most significantly differentially regulated GO pathways, revealing that SAS3-transduced mice show significant up-regulation of pathways associated with inflammation and immune response. **(B)** Immunostaining for GFAP and IBA1 in the hippocampus, showing activated microglia clustered around SAS-induced aggregates. Scale bar equals 50 μm.

